# Marine spatial planning to enhance coral adaptive potential

**DOI:** 10.1101/2024.08.27.609972

**Authors:** Daniel L. Forrest, Lisa C. McManus, Eden W. Tekwa, Daniel E. Schindler, Madhavi A. Colton, Michael M. Webster, Helen E. Fox, Timothy E. Essington, Stephen R. Palumbi, Peter J. Mumby, Lukas DeFilippo, Steven R. Schill, F. Joseph Pollock, Malin L. Pinsky

## Abstract

Ocean warming interacts with local stressors to negatively affect coral reefs. The adaptive capacity of reefs to survive these stressors is driven by ecological and evolutionary processes occurring at multiple spatial scales. Marine protected area (MPA) networks are one solution that can address both local and regional threats, yet the impacts of MPA network design on adaptive processes remains unclear. In this paper, we used an eco-evolutionary model to simulate hypothetical MPA configurations in the Caribbean, Southwest Pacific and Coral Triangle under projected warming. We found that protecting thermal refugia (i.e., cooler reefs) largely benefited corals inside the refugia while other reefs declined. In contrast, protecting a diverse habitat portfolio led to increased coral cover both inside and outside of the MPA network. We then quantified the thermal habitat and connectivity representations of reefs both inside and outside existing MPA networks across each region. Most strikingly, reefs in current MPA networks in the Southwest Pacific and Coral Triangle are approximately 2 °C cooler than reefs outside the MPA networks, while the Caribbean’s MPA network is approximately 1 °C warmer than reefs outside the network, based on mean temperatures from 2008-2018. These results suggest that the Caribbean MPA network is poised to protect sources of warm-adapted larvae but not destinations, and the opposite is true of the Southwest Pacific and Coral Triangle. Our results suggest that 1) by protecting sites with particular temperature and connectivity characteristics, marine spatial planning may alter eco-evolutionary processes to enhance or inhibit the adaptive capacity of a reef network and 2) the distribution, extent, and effectiveness of local interventions have the potential to affect regional distributions of coral cover beyond what would be expected from local benefits alone, due to the potentially wide-reaching effects of larval dispersal and gene flow.

## Introduction

The increasing frequency and severity of marine heat waves are predicted to result in global scale coral reef declines by the mid 21st century (van Hooidonk et al. 2013; Hoegh-Guldberg et al. 2017; Logan et al. 2021; Setter et al. 2022), highlighting the importance of reducing emissions for the persistence of corals (IPCC 2021). However, while corals are primarily threatened by global stressors, management and conservation actions to protect reefs tend to focus on relieving reefs of local stressors (White & Vogt 2000; Aswani et al. 2015). Effective local management—including interventions that improve watershed health (Fabricius 2005; Wooldridge & Done 2009; De’ath et al. 2012; Rodgers et al. 2012) and reduce fishing of herbivores (Mumby et al. 2007; Burkepile & Hay 2008; Suchley & Alvarez-Filip 2017)—can help coral reefs resist and recover from the detrimental impacts of climate change (Donovan et al. 2021; Gove et al. 2023). To that end, marine protected areas (MPAs), where extractive activities such as fishing are restricted (Sala & Giakoumi 2018), are a common tool to reduce local impacts on marine systems (Selig & Bruno 2010; Halpern et al. 2010). Due to the interconnectedness of marine habitats, implementing groups of MPAs—an MPA network—can bridge the gap from local- to regional-scale conservation (Ovando et al. 2021; Colton et al. 2022).

MPA networks attempt to maintain larval connectivity among reefs, leading to significant ecological benefits (Gaines et al. 2010; Balbar & Metaxas 2019; Harrison et al. 2020). In particular, theoretical studies have highlighted the role of dispersal stepping stones that connect sites across the network (Treml et al. 2008; Kininmonth et al. 2011; Watson et al. 2011) and the importance of protecting sites that act as both demographic sources and sinks for network persistence (Jacobi & Jonsson 2011; Krueck et al. 2017; Kininmonth et al. 2019). Additionally, incorporating connectivity metrics into conservation planning software (e.g., Marxan) can effectively account for connectivity patterns (Beger et al. 2010, 2015; Schill et al. 2015; Krueck et al. 2017; Tong et al. 2021), resulting in higher biomass and yields in the case of fisheries (White et al. 2014). While recent reef prioritization frameworks explicitly call for the consideration of larval connectivity metrics in constructing MPA networks (Beger et al. 2015; Beyer et al. 2018), effectively linking connectivity science and MPA management remains an ongoing challenge in marine conservation (Lagabrielle et al. 2014).

Other important considerations for MPA networks are eco-evolutionary feedbacks, which occur because ecological (e.g., competition, dispersal) and evolutionary (e.g., local selection, gene flow) processes interact (Post & Palkovacs 2009; Govaert et al. 2019). Under certain conditions, adaptation as a result of these feedbacks—eco-evolutionary adaptation—is expected to mitigate coral cover loss during ocean warming and facilitate recovery post-warming (Bay et al. 2017; Matz et al. 2018, 2020; Logan et al. 2021; McManus et al. 2021b). One type of feedback occurs when competition (an ecological process) affects natural selection (an evolutionary process) to change the community composition and phenotypes of populations, which then impacts ecological dynamics (Post & Palkovacs 2009; Henry 2017). Another type of feedback occurs through dispersal, which impacts the spread of adaptive and non-adaptive alleles throughout the network, affecting local demographics (Matz et al. 2018, 2020; McManus et al. 2021b). Due to these feedbacks and environmental heterogeneity across the network, conservation outcomes from localized reef protection may run counter to expectations in networks (McManus et al. 2021a), potentially diminishing the effectiveness of conservation plans.

Guidelines for eco-evolutionary MPA network design suggest that protecting a diversity of environmental conditions can favor adaptation (Walsworth et al. 2019), partly because habitat representation of thermal regimes may be critical for maintaining the standing genetic variation of a coral reef network (Howells et al. 2013; Baums et al. 2019). However, few studies have explored the performance of MPA network designs on realistic seascapes (Mumby et al. 2011; Tong et al. 2021; Ovando et al. 2021), such that the effects of MPA network configuration on reef persistence remain unclear. Furthermore, while we understand which network attributes lead to persistence, there has been little to no attempt to apply those concepts to existing protection networks.

Here, we aimed to explore and generate new hypotheses regarding MPA network effects on regional coral reef eco-evolutionary dynamics. We modeled the effects of alternative MPA network configurations in the context of thermal environments and adaptive capacity across the Caribbean, Southwest Pacific, and Coral Triangle. By utilizing realistic seascapes in our simulations, we sought to generate broader hypotheses about spatial design principles for MPA networks, rather than prescribing protection for specific sites. This approach allowed us to explore general patterns in how network design may influence eco-evolutionary processes and coral persistence under climate change scenarios. We then analyzed characteristics of existing MPA networks in each region to evaluate whether existing MPA networks are well-configured to enable eco-evolutionary processes that lead to coral persistence.

## Materials and Methods

### Eco-evolutionary model

To model the efficacy of different MPA network designs, we modified an existing eco-evolutionary model (McManus et al. 2021a) that has been applied to regional-scale coral reef networks under climate change (McManus et al. 2021b) and restoration scenarios (DeFilippo et al., 2022). We modeled the population dynamics and trait evolution of two spawning coral types: one that grows rapidly and has a narrower thermal tolerance, and one that grows slowly and has a wider thermal tolerance (Darling et al., 2012). On each reef, we tracked the change in cover of each coral type and their mean trait value, the mean thermal optimum. Coral cover responded to population growth, competition and larval immigration. The thermal optimum of each type evolved due to gene flow among sites and stabilizing selection in response to changing sea surface temperatures.

We also modeled macroalgae, which competed with coral for space (Mumby et al., 2007). Macroalgae were included in the framework to serve as a competitor for corals and to facilitate the implementation of MPAs. As such, they did not experience evolutionary effects nor dispersal among patches. Furthermore, because macroalgae tend to be much less sensitive to temperature than coral, macroalgal population dynamics were assumed to be insensitive to temperature (Anton et al. 2020).When a reef was designated as an MPA, macroalgal mortality was locally increased as a proxy for the effects of common management interventions, such as reduced fishing pressure on herbivores that graze on macroalgae or a reduction of land-based nutrient inputs that favor macroalgal growth. We note that our definition of an MPA in the model is not specific to ‘no-take’ marine reserves that prohibit all fishing and other extractive activities (Sala & Giakoumi 2018). Rather, an MPA designation in the model denotes the reduction of local stressors that promote macroalgae growth. Full details of the eco-evolutionary framework are presented in Appendix S1.

### Simulations

We applied the model to simulate coral dynamics over a hindcast period of 149 years (1870-2018) without MPAs and a projection period with MPA network implementation of 282 years (2018-2300). These time periods represented the full temporal coverage of the sea surface temperature (SST) datasets from HadISST1 SST (Rayner 2003) and GISS E2 H (Schmidt et al. 2014) for the hindcast and forecast periods, respectively. Initial cover of fast coral, slow coral, and macroalgae, and coral mean trait values for the hindcast were obtained from the end of a 500-year burn-in period. Burn-in simulations were implemented with constant temperature from 1870 and without larval dispersal. Parameters were set to facilitate the coexistence of the fast coral, slow coral, and macroalgae within empirically observed ranges when no MPAs were implemented (Tekwa et al. 2021). Reef area, potential connectivity and temperature were the only inputs that varied across reefs and regions, while all other parameters were held constant (Appendix S2). This approach supported the comparison of MPA strategy effectiveness both within and among regions. Note that the parameters used here have been thoroughly explored in our previous studies (McManus et al., 2021a; McManus et al., 2021b). These prior analyses have demonstrated the robustness of the model across a range of parameter values, providing a solid foundation for its application in the current study.

Potential connectivity data from previously published larval connectivity matrices were incorporated in the simulations. In short, these matrices were generated by combining ocean circulation models with biophysical larval tracking in the Caribbean (Schill et al. 2015), Coral Triangle (Thompson et al. 2018) and Southwest Pacific (Treml et al. 2008) regions. All three matrices were developed to approximate the passive dispersal of a coral-like species among reefs (see Appendix S3 for details regarding connectivity data). The biophysical models do not simulate on-reef dispersal processes in fine detail, since these fine-scale processes are not computationally tractable at large spatial extents. The number of reef units (Caribbean: 423 reefs, Southwest Pacific: 583 reefs, Coral Triangle: 2083 reefs) and reef locations were also obtained from these outputs. The ocean circulation models have been tested against observations (Treml et al. 2008; Schill et al. 2015; Thompson et al. 2018), and while the resulting larval connectivity matrices are untested, there is evidence that oceanographic distance from biophysical simulations predicts genetic connectivity, especially at large spatial scales (Davies et al. 2015; Riginos et al. 2019; Galaska et al. 2021; Wang et al. 2022; Fitz et al. 2023).

The hindcast (HadISST1) and projection (GISS E2 H) SST time series datasets both had a resolution of 1×1 degree latitude x longitude. We applied the delta method (Hay et al. 2000; Fowler et al. 2007; Ramirez Villejas & Jarvis 2010) to downscale these products using a climatology from NOAA OISST V2 from 1982-2010 which had a higher resolution of 0.25×0.25 degree (Reynolds et al. 2007). We generated SST trajectories for the RCP 4.5 climate scenario (Pachauri et al. 2015) for each reef within the three regions. This scenario represents a reduction of greenhouse gas emissions and a subsequent stabilization of temperatures after a global mean rise between 2 and 3°C (Pachauri et al. 2015).

### MPA prioritization

We conducted simulations in which MPAs represented 0%, 10%, 20%, 30% and 50% of reefs in the network. We focused our main results on the 30% coverage scenario, as this is a common marine conservation target (Klein et al. 2010; IUCN 2021; Kalinina 2021). The primary simulations with 30% coverage consisted of 126 MPA sites and 297 non-MPA sites in the Caribbean, 174 MPA sites and 409 non-MPA sites in the Southwest Pacific, and 624 MPA sites and 1459 non-MPA sites in the Coral Triangle. We tested the following nine configurations for MPA prioritization (Table 1): 1. ‘cold,’ 2. ‘hot,’ 3. ‘even temperature,’ 4. ‘eigenvector centrality,’ 5. ‘isolation,’ 6. ‘preadapted larvae,’ 7. ‘pr05’, 8. ‘evenly spaced’, and 9. ‘representative.’ Broadly, MPA configurations were based on temperature (1-3), connectivity (4 & 5), a combination of temperature and connectivity (6 & 7), or site heterogeneity (8 & 9) (Fig. 1).

**Figure 1.**
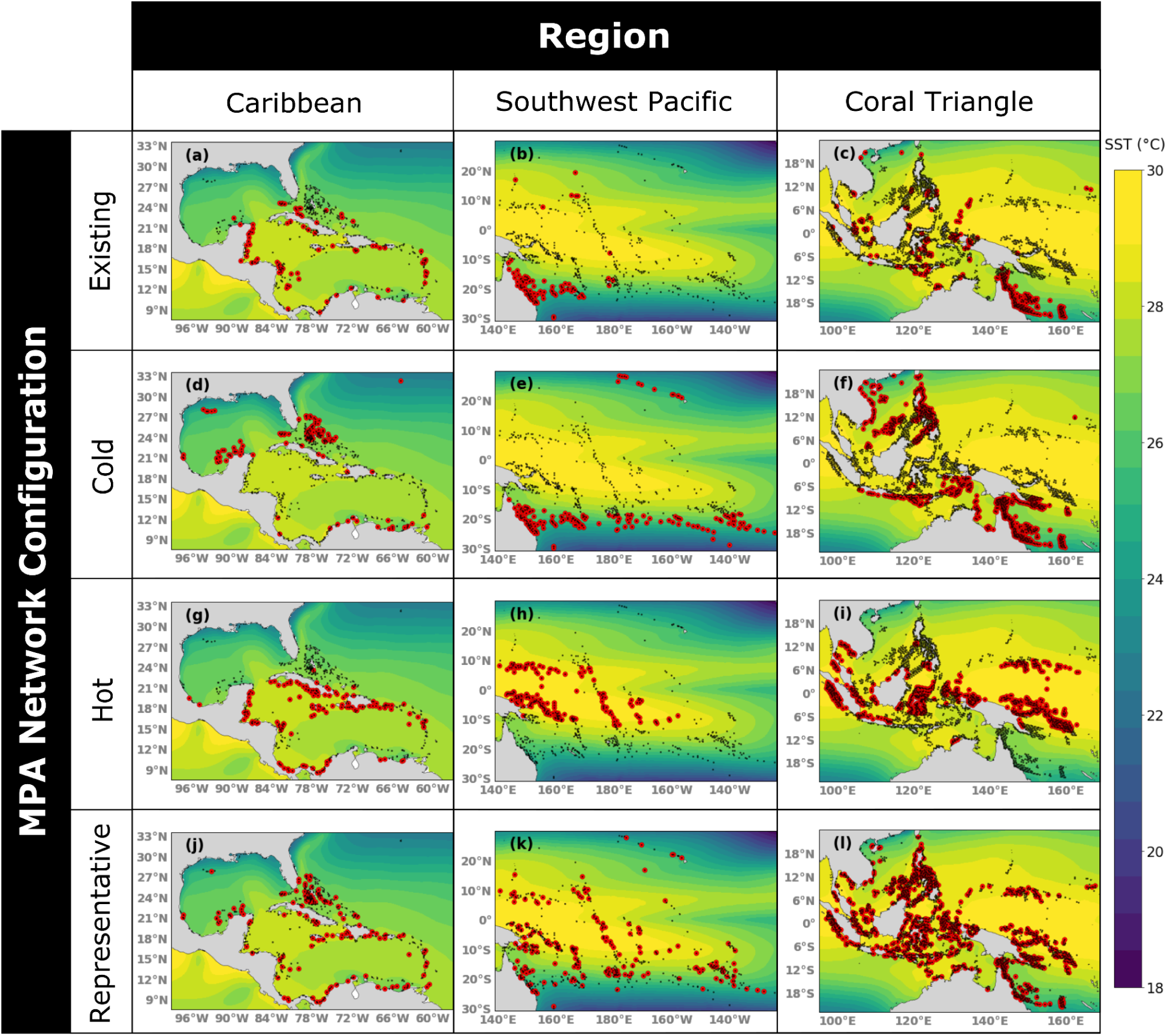
Maps of reef sites designated as marine protected areas (MPAs) under four different network configurations: Existing MPAs (a-c), Cold (d-f), Hot (g-i), and Representative (j-l). Black dots represent all individual reef patches, red highlights indicate an MPA designation, and the heatmap background represents mean SST based on the high resolution NOAA OISST V2 output from 1982-2010 (0.25 x 0.25 degree; Reynolds et al. 2007). The representative configuration displayed is one of 10 that were simulated.

**Table 1.** Descriptions of site-selection strategies for marine protected areas.

| Category | Strategy | Algorithm |
| --- | --- | --- |
| Temperature | 1. Cold | Select the x% coldest reefs* |
|  | 2. Hot | Select the x% hottest reefs* |
|  | 3. Even temperature | Select reefs sorted by temperature at a regular interval to reach x%* |
| Connectivity | 4. Eigenvector centrality | Select the x% of reefs with the highest eigenvector centrality (i.e., reefs that are important to metapopulation growth) (Ovaskainen and Hanski, 2003) |
|  | 5. Isolation | Select the x% of reefs with the lowest destination strength (i.e., reefs that had the lowest potential for gene flow to support evolutionary adaptation [Lenormand, 2002]) |
| Temperature & connectivity | 6. Preadapted larvae | Select the x% of reefs with the strongest gene flow from warmer sites* |
|  | 7. pr05 | Select the x% of reefs with the highest proportion of site connections that are at least 0.5 °C warmer* (Matz et al., 2020) |
| Heterogeneity | 8. Evenly spaced | Select the x% of reefs such that they are evenly spread across the network based on geographic location |
|  | 9. Representative | Select x% at random to capture a representative subset of the region (i.e., reefs that approximate a portfolio that captures network-wide variation of temperature and connectivity characteristics [Figge, 2004; Schindler et al., 2015]) |

\*Indicates that temperatures for calculation were based on the 2008-2018 climatology.

The three temperature configurations involved sorting reefs based on their mean climatology from 2008 to 2018 (the last ten years of the hindcast) and then selecting either the coldest sites (‘cold’), hottest sites (‘hot’), or sites at regular intervals (sorted by SST) depending on the percentage of network protection (‘even temperature’). Colder sites represent one definition of climate refugia, or sites that are likely to maintain habitable conditions under environmental change (Keppel et al. 2015), while warmer sites are expected to be sources of warm-adapted larvae (Norberg et al. 2012).

The ‘eigenvector centrality’ configuration selected those reefs that contribute the most to the metapopulation growth rate of the network and was calculated based on the left eigenvector associated with the leading eigenvalue of the connectivity matrix (Ovaskainen & Hanski 2003). For ‘isolation’, we selected sites that had the lowest destination strength scores. Destination strength was the sum of all potential immigrating larval connections based on each region’s simulated larval connectivity matrix (Thompson et al. 2018). Destination strength was calculated as *S_a_* = Σ*_b_*D*_ab_*, where *S_a_* was the destination strength at site *a* and *D_ab_* was the potential connection strength from site *b* to *a* (from the connectivity matrix).

For ‘preadapted larvae,’ we summed the average differences in SST among all sites and each focal site across years 2008 to 2018, scaled by connectivity strength and the reef areas of both destination and source reefs. We then selected the sites that were most strongly connected to warmer reefs. For ‘pr05,’ the procedure was identical to that for ‘preadapted larvae’, except that source sites had to be at least 0.5 °C warmer to be included in the calculation of this metric for each reef (Matz et al. 2020).

For ‘evenly spaced,’ we first chose the site with the highest eigenvector centrality score. We then selected the next site that was furthest from it based on geographic distance, then selected the next site that maximized the distance from the other two locations, and so on until the appropriate number of sites was selected.

For ‘representative,’ we randomly selected sites to capture a representative subset of the region. We then repeated this ten times to create ten independent networks to compare. These selected reefs approximated a portfolio that captured network-wide variation of temperature and connectivity characteristics (Figge 2004; Schindler et al. 2015). All configurations and their implementation are described in Table 1.

### Analysis of existing MPA network characteristics

To understand whether existing MPA networks can support eco-evolutionary adaptation, particularly through environmental diversity, we quantified thermal habitat and larval connectivity across MPA networks in the Caribbean, Southwest Pacific, and Coral Triangle. Specifically, we assessed whether the distributions of temperature and connectivity metrics in each region’s MPA network were representative of the broader region. We obtained MPA locations from Protected Planet (UNEP-WCMC & IUCN 2020). For each region, all sites that fell within MPA polygons from Protected Planet were designated as MPAs in our model. We used the connectivity, reef area, and temperature data as described above.

We compared MPA and non-MPA subsets of our network for the following seven connectivity and temperature characteristics: 1. initial sea surface temperature (*T_0_*; mean temperature from 2008-2018), 2. the change in sea surface temperature over the full climate change projection (*ΔT*; [mean SST from 2290-2300] - [mean SST from 2008-2018]), 3. destination strength (*S*), 4. local retention (*L*), 5. self-recruitment (*R*), 6. the proportion of incoming connections from reefs that are at least 0.5 °C warmer relative to the local reef temperature (pr05 sensu (Matz et al. 2020)), and 7. reef area (Area; reef area in m^2^). We considered *T_0_* because cooler reefs can potentially benefit from the existence of warmer-adapted larvae in the network (Norberg et al. 2012); this latter mechanism is more explicitly quantified by the pr05 metric. The amount of thermal stress a reef was projected to experience was captured in *ΔT*: lower values implied less future mortality due to bleaching (Hoegh-Guldberg et al. 2017) and a higher chance of evolutionary adaptation (Lindsey et al. 2013). Destination strength was calculated as described above and local retention was the probability of self-connection relative to the sum of all outgoing connections: 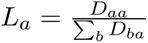. Self-recruitment was the probability of self-connection relative to the sum of all incoming connections: 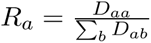. High destination strength indicates strong larval input rates to facilitate demographic rescue (Brown & Kodric-Brown 1977), while high local retention suggests long-term persistence (Botsford et al. 2009). High self-recruitment is associated with low immigration and high potential for local adaptation (Lenormand 2002). We obtained areas for each region from the connectivity matrix sources (Appendices S3 and S4). Because these analyses were meant to provide a description of observed differences (and not to make inferences about a larger population of protected and unprotected networks), and because the assumption of statistical independence was violated twice–due to comparing all MPA and non-MPA sites from the same region, and due to the potential spatial autocorrelation of MPA designations–we did not perform statistical analyses to compare these metrics across MPA and non-MPA reefs. All code and simulated data are available from Zenodo at https://doi.org/10.5281/zenodo.8432964.

## Results

### MPA configuration simulations

We focus our presentation on hot, cold, and representative MPA configurations because these produced the most qualitatively different results and represent end-points on a continuum of active debate in marine conservation. Regardless of network configuration, higher proportions of MPAs in the seascape led to higher coral cover. This suggests that the primary effect of MPAs was a network-wide demographic boost associated with increased macroalgal mortality (which decreased competition with coral) at MPA sites (Figs. 2a-c, Appendices S5 and S6). Mean coral cover declined in MPA network configurations during the period of rapid temperature increase, from approximately 2018-2050, followed by a recovery period starting at ∼2100 (Fig. 2a,b,c and Appendix S7). Mean coral cover without MPA implementation followed similar qualitative patterns in terms of decline and recovery in all three regions. However, minimum coral cover, occurring at ∼2100, differed between scenarios with and without MPAs: compared to the scenario without MPAs, configurations with 30% MPA coverage increased minimum cover by factors of 1.84 to 2.26 in the Caribbean, 1.18 to 1.21 in the Southwest Pacific and 1.16 to 1.23 in the Coral Triangle. At the local scale, sites designated as MPAs generally had higher cover relative to the scenario without MPAs (Appendix S8).

**Figure 2.**
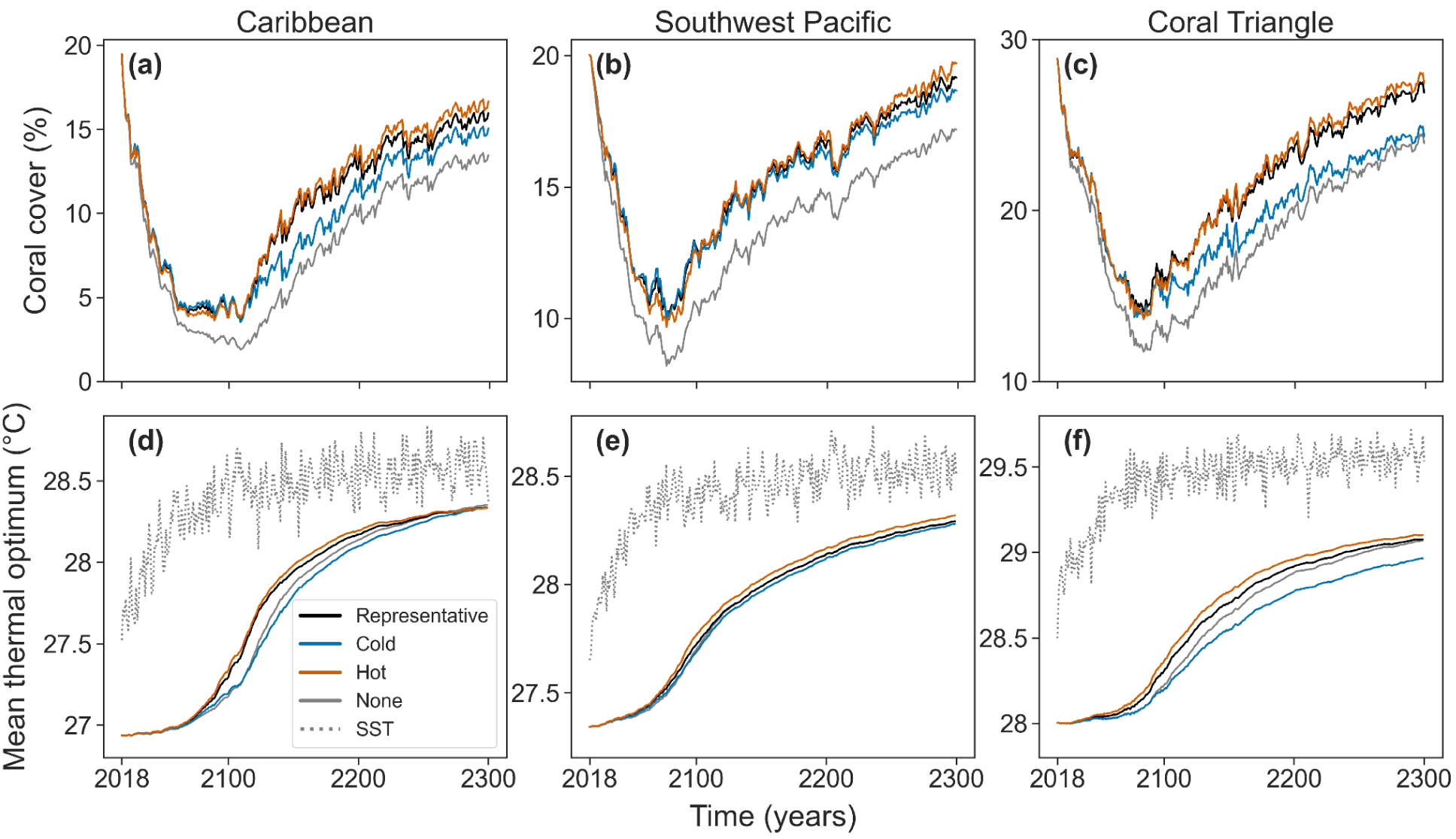
Trajectories of mean total coral cover (a, b, c) and mean thermal optimum (d,e,f) for the representative (solid black line), cold (blue line), and hot (orange line) MPA network configurations and the scenario with no MPAs implemented (solid gray line) across the three study regions. Note that y-axes vary by plot to highlight differences among configurations.

There were broad similarities across the three regions in the relative performance of alternative MPA strategies. The dynamics of the mean trait value revealed that the hot and representative configurations allowed the corals to better evolve higher thermal tolerance through time (Fig. 2d-f, Appendix S9). Across all three regions, the cold configuration was least able to facilitate adaptation to increasing temperatures. This pattern becomes most clear after 2100 for the Caribbean and Coral Triangle and after 2250 in the Southwest Pacific (Fig. 2a-c). Initially and through the decline period, the cold strategy maintained the highest coral cover but only by a small magnitude relative to other strategies. During the recovery period, the hot strategy led to the highest coral cover in all three regions, followed closely by the representative strategy, which in turn outperformed the cold strategy by 2.7 to 10% in cover across regions (Appendices S10-S13).

Site-selection strategies had different effects on coral populations inside and outside of MPAs. In general, the cold configuration led to higher coral cover inside MPAs relative to outside and the hot configuration led to higher coral cover outside MPAs relative to inside. The representative strategy led to cover that was intermediate relative to the cold and hot configurations and was more evenly distributed among MPA and non-MPA reefs (Fig. 3 and Appendix S14). An exception to this trend was in the Caribbean network, where the hot strategy had higher coral cover inside MPAs compared to other regions (though not as much as the cold strategy). This result can likely be attributed to the coincidentally higher destination strength in the Caribbean (Appendix S15 and S16). To quantify the relative association of initial sea surface temperature, destination strength and MPA status on minimum coral cover, we fit multiple regression models for each iteration of a representative MPA configuration for all three regions. In general, minimum coral cover was negatively correlated with initial sea surface temperature, positively correlated with destination strength, and sites selected as MPAs experienced higher minimum cover (i.e., a higher intercept). In the Southwest Pacific and Coral Triangle, initial SST had the largest magnitude coefficient, followed by destination strength and MPA status. In the Caribbean, DS had the largest magnitude, followed by initial SST. For all three regions, MPA status had the lowest magnitude coefficients (Appendices S17-S19).

**Figure 3.**
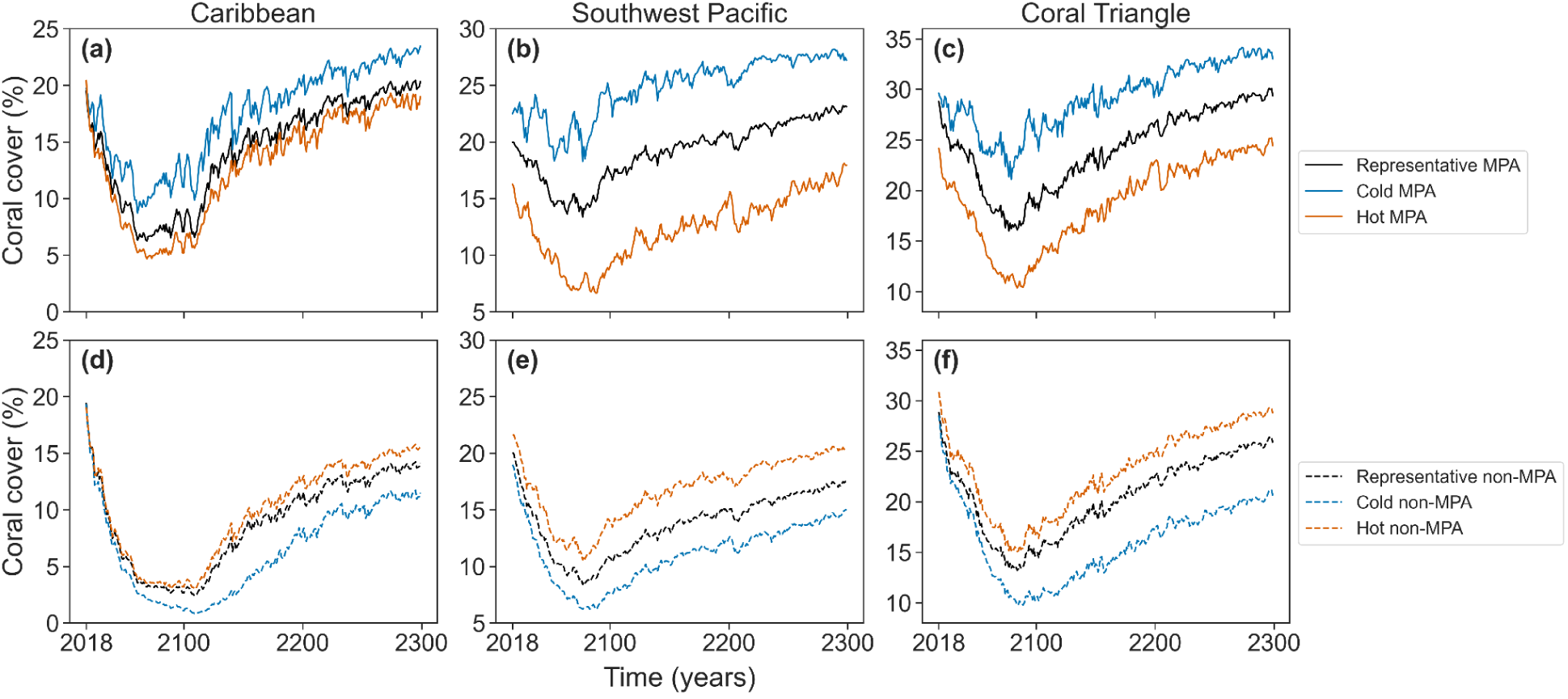
Trajectories of mean cover in MPA sites (a-c, solid lines) and non-MPA sites (d-f, dashed lines) for the representative (black), cold (blue) and hot (orange) configurations in the three regions.

Strikingly, placing MPAs on cold sites produced worse outcomes for non-MPA reefs (i.e., lower coral cover) relative to the scenario without any MPAs (Appendix S20). The effect likely occurred because protecting cold sites favored maladaptation of corals under rising temperatures, which more than offset the lower competition from macroalgae that occurred with MPA implementation (Appendix S9).

To better understand differences among configurations, we also calculated coral cover relative to the representative configuration at the network level, within MPA sites, and at non-MPA sites (Fig. 4). Figure 4 aims to disentangle whether the effect of MPAs on the distribution of coral cover was due to the underlying characteristics of the selected sites or the added eco-evolutionary effect of the MPA network configuration. In other words, we ask: how much coral cover would exist on the same sites when MPAs are implemented on the coldest or hottest 30% of the network relative to when MPAs are implemented with the representative configuration? In the Caribbean and Southwest Pacific, the hot configuration led to lower coral cover relative to the representative configuration at the beginning of the simulation, while the cold configuration showed the opposite trend; this pattern was reversed starting from ∼2100 (Fig. 4a,b). In the Coral Triangle, both cold and hot configurations had approximately equivalent or lower coral cover relative to representative. In this region, the cold and hot strategies diverged at ∼2100, and from ∼2150 onwards the hot and cold configurations led to higher and lower cover relative to representative, respectively (Fig. 4c). Relative to the representative strategy, non-MPA sites in all three regions had lower coral cover with the cold configuration and higher cover with the hot configuration; this pattern was flipped for MPA sites.

**Figure 4.**
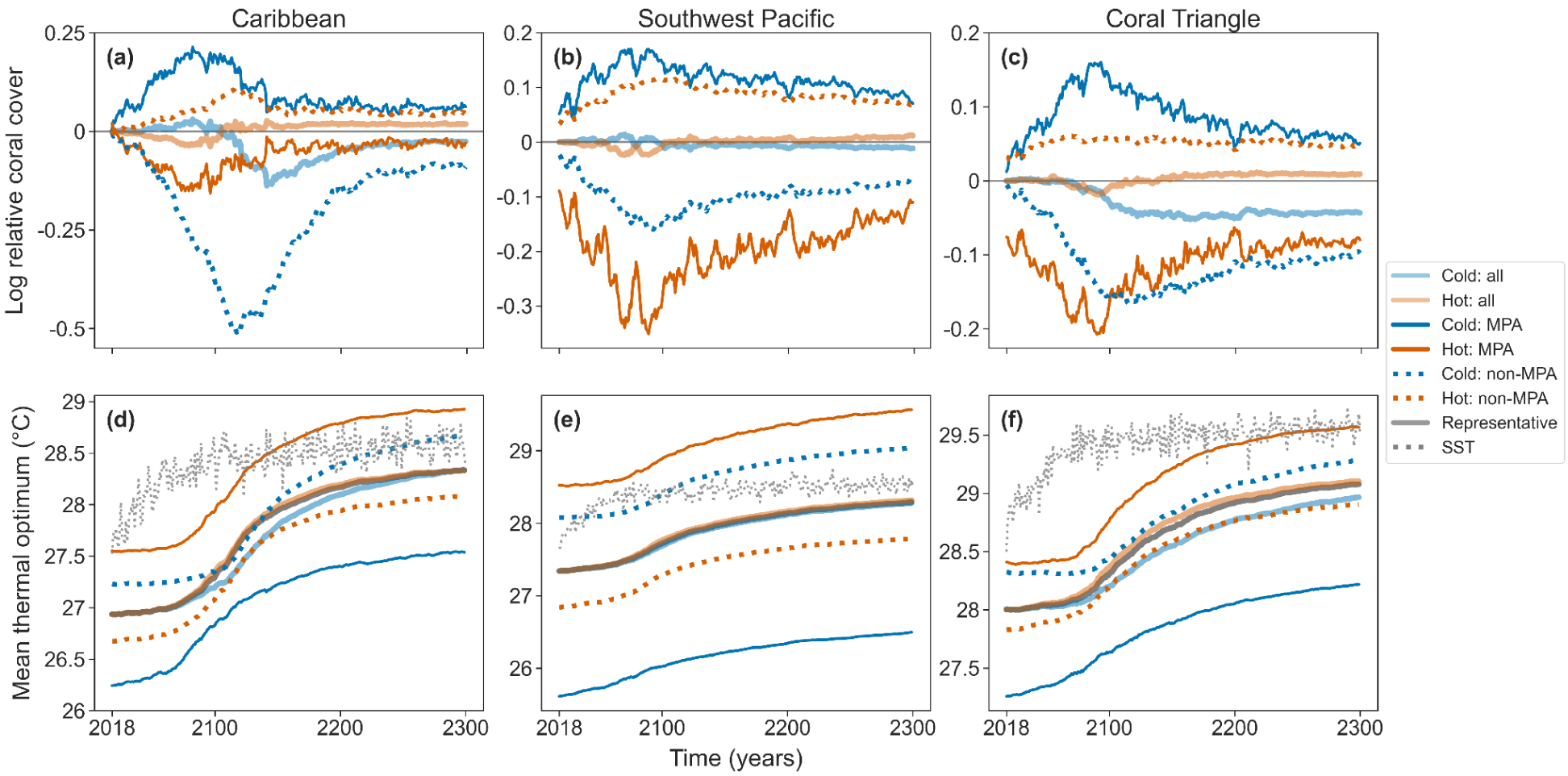
Relative coral cover calculated as log_10_(mean coral cover in strategy X / mean coral cover in representative strategy) for cold and hot configurations (a-c) and corresponding mean thermal optima (d-f) calculated for all sites, MPA sites and non-MPA sites in the Caribbean, Southwest Pacific and Coral Triangle.

Some MPA configurations led to counter-intuitive results whereby coral cover was greater outside of MPAs than inside (Appendix S14). In the Caribbean and Southwest Pacific, the isolation strategy—protecting sites with the lowest potential for external recruitment and gene flow—led to higher cover outside MPAs at the beginning of the simulation and led to higher cover inside MPAs during the recovery period and onwards (Appendix S14d,e). Interestingly, the isolation configuration in the Coral Triangle led to higher coral cover outside MPAs for the duration of the simulation (Appendix S14f). Finally, the evenly spaced (selecting sites that are evenly distributed across space), even temperature (selecting sites across the entire temperature gradient), pr05, and preadapted larvae strategies all supported higher coral cover in MPA sites than non-MPA sites across regions (Appendix S14g-i).

Trajectories exhibited by each coral type largely echoed those for the combined coral community but also revealed nuanced patterns. Fast coral declined by ∼2075 and began recovery by ∼2100, whereas slow coral declined more slowly and less, but only experienced marginal recovery over the full simulation. Thus, the community shifted from slow- to fast-coral dominance. By 2300, the mean thermal optimum for fast coral was within the range of interannual SST variation, while the mean optimum for slow coral lagged by approximately 1-1.5 °C (Appendix S21). We can likely attribute this difference to the thermal tolerance breadth: slow corals had a broader tolerance and experienced lower selective pressure that resulted in a slower rate of adaptation.

Under the cold configuration, slow coral cover in non-MPA Caribbean sites declined to near zero, indicating that those sites were primarily composed of fast coral. Fast coral in the Southwest Pacific and Coral Triangle experienced a larger difference in cover between MPA and non-MPA reefs under the representative configuration (higher in MPAs) than in the hot configuration (lower in MPAs and relatively even in the Southwest Pacific and Coral Triangle, respectively) (Appendix S22).

### Characteristics of existing MPAs

Our gap analysis showed that MPAs in the Caribbean are located in generally warm sites, with much less protection in cold sites for the region (Fig. 5). The opposite was true for the Southwest Pacific and the Coral Triangle, where MPA sites have been placed in generally cooler sites and are underrepresented in warm sites. Variation in the thermal profiles of protected sites can be measured as the standard deviations of initial SST (mean 2008-2018) within regions. In the Caribbean, MPA sites (n=81, 19%) had a mean ± standard deviation SST of 28 ± 0.62 °C, while non-MPA sites (n=342, 81%) had a mean of 27 ± 0.80 °C. In the Southwest Pacific, MPA sites (n=92, 16%) had a mean SST of 26 ± 1.2 °C while non-MPA sites (n=491, 84%) had a mean of 28 ± 1.4 °C. MPA sites in the Coral Triangle (n=291, 14%) had a mean SST of 27 ± 1.3 °C while non-MPA sites (n=1792, 86%) had a mean of 29 ± 0.61 °C (Appendices S23-S25).

**Figure 5.**
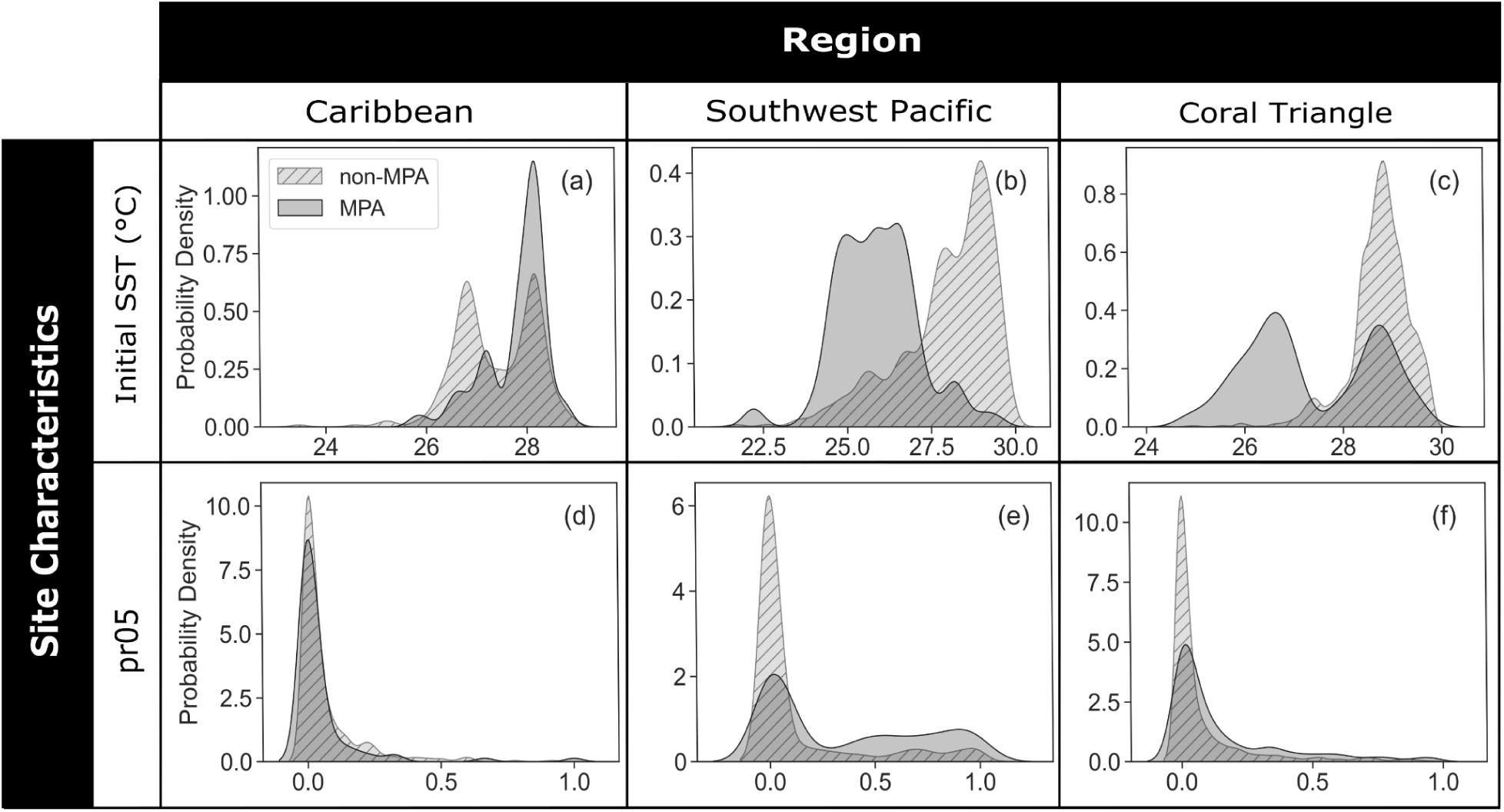
Site characteristics in existing MPA vs. non-MPA sites. Kernel density functions of initial sea surface temperature (T_0_; °C) (a-c) and pr05 (d-f) across the three study regions (columns).

MPA and non-MPA sites differed in their pr05 values, which quantifies the proportion of site connections that are at least 0.5 °C warmer, among all regions (Fig. 4g-l). Caribbean MPAs were located in sites with somewhat lower pr05 (0.05 ± 0.14) than non-MPA sites (0.07 ± 0.12). In the Southwest Pacific and Coral Triangle, MPAs have been placed in sites with exceptionally high pr05 (Southwest Pacific: MPA = 0.36 ± 0.37, non-MPA = 0.13 ± 0.27; Coral Triangle: MPA = 0.15 ± 0.23, non-MPA = 0.08 ± 0.17). These regional differences in pr05 of the MPAs aligned with the regional differences in site temperatures of the MPAs. In other words, sites with high pr05 tend to be cooler since those sites are more likely to experience warmer connections. Other metrics revealed much smaller differences among MPA and non-MPA sites (Appendices S23-S26).

## Discussion

Mounting evidence from eco-evolutionary models indicates that marine connectivity and evolution can affect reef resilience under ocean warming (Bay et al. 2017; Walsworth et al. 2019; Matz et al. 2020; Logan et al. 2021; McManus et al. 2021b). Yet, while coral conservation planning has considered a wide range of factors in site selection recommendations, it has not typically considered eco-evolutionary dynamics (Beyer et al. 2018; McClanahan et al. 2023). The few studies that have considered eco-evolutionary dynamics in this context used network optimization approaches (e.g., Beger et al., 2014b; Mumby et al., 2011), and did not explicitly link these hypothesized networks to impacts on reef- and regional-scale coral cover. Here we examined three seascapes, finding that MPA networks from all three regions currently encompass a skewed temperature distribution compared to the full region. We found that MPA configuration performance, as assessed with maximum coral cover, varied temporally (pre- vs. post-temperature stabilization) and spatially (e.g., enhanced coral cover inside vs. outside the MPA network). Overall our simulation results suggested that 1) by protecting sites with particular temperature and connectivity characteristics, marine spatial planning may alter eco-evolutionary processes to enhance or inhibit the adaptive capacity of a reef network and 2) the distribution, extent, and effectiveness of local interventions have the potential to affect regional distributions of coral cover beyond what would be expected from local benefits alone, due to the potentially wide-reaching effects of larval dispersal and gene flow.

### Eco-evolutionary simulations

MPA configurations substantially differed in the proportion of coral cover maintained inside and outside of the MPA network. Differences in cover over time can be attributed to the temperature and connectivity characteristics of protected reefs, as all other model parameters were identical among sites. In general, protecting the coldest reefs in the network maintained more coral cover inside the MPA network, protecting the hottest reefs maintained more coral cover outside of the MPA network, and protecting diverse habitats (representative configuration) maintained comparatively even coral cover within and outside the MPA network. To a lesser extent, design strategies differed in the amount of coral supported before (<2100) and after temperature stabilization (>2100). The cold configuration was better at mitigating near-term declines, but did not achieve as complete a recovery by 2300. Coral cover in the hot configuration declined most in the near-term, but supported the most substantial long-term recoveries. Finally, the representative configuration best maintained cover across all time periods.

The representative configuration best maintained reef health through the short and long term and throughout the entire region, supporting previous findings that protecting diverse habitats facilitates multiple adaptation pathways: local adaptation, demographic rescue, and beneficial gene flow (Walsworth et al. 2019; Colton et al. 2022). We extend this work by modeling coral dynamics on spatially realistic seascapes and exploring the relative impacts of MPA implementation to individual reefs inside and outside of MPAs. When only sources *or* destinations of warm-adapted larvae are protected, the effectiveness of one or more potential adaptation pathways is reduced (McManus et al. 2021b). Surprisingly, the isolation configuration led to higher cover at non-MPA sites for at least the beginning of the simulations. This may be driven by the underlying characteristics of MPAs selected in this configuration (Appendix S27), but in general, this suggests that selecting sites based on a single metric across a region may lead to counterintuitive outcomes due to the complexity of real seascapes. We also note that selecting sites with the express goal of representing the temperature variation of a region (even temperature configuration) resulted in higher cover at MPA sites relative to non-MPA sites. This implies that the approximately even coral cover within and outside of MPAs observed in the representative configuration was due to selecting sites that represented a diversity of temperature *and* connectivity characteristics.

These results lend support to the application of portfolio theory (Figge 2004; Schindler et al. 2015) in conservation planning. Representative and diversity-based MPA network design strategies, a habitat-based application of portfolio theory, were once popular in coral reef conservation and management (Fernandes et al. 2005). However, most research over the last decade has proposed refugia-based strategies with a narrow definition, i.e., reefs that avoid the greatest disturbance (Maina et al. 2011; Beyer et al. 2018; Wilson et al. 2020; van Woesik et al. 2022), while some researchers are proposing to broaden the definition of refugia to include considerations of coral community composition and warming response (McClanahan et al. 2023). Our results indicate that the representative configuration, which provided protection to reefs with diverse temperature and connectivity profiles, supported both sources and destinations of warm-adapted larvae. In contrast, the cold configuration, approximating a refugia approach, primarily retained corals within the MPA network. This suggests that refugia-based strategies (as determined by reef temperature) may not benefit reefs and associated human communities in locations outside of the MPA networks as much as those within MPAs, whereas representative strategies would likely provide coral cover benefits to reefs within and outside of the MPA network. These results warrant additional research and could serve as hypotheses for future empirical research that aims to investigate the viability of portfolio and other habitat-diversity approaches to coral reef conservation.

These collective findings are likely due to the eco-evolutionary feedbacks driven by the movement of individuals (e.g., coral larval dispersal) and local adaptation (Lenormand 2002; Govaert et al. 2019), including the degree to which the MPA networks favor maladaptation (cold configuration), preadaptation (hot), or diverse genotypes (representative). We demonstrate that MPA network configurations can influence these underlying feedbacks and lead to complex outcomes for coral persistence. In a metapopulation, persistence at a given location can be supported by the immigration of individuals that are preadapted to impending environmental stress (Bell and Gonzalez, 2011). Under increasing thermal stress, cooler sites largely rely on preadapted input from warmer sites, while warmer sites primarily rely on local adaptation (Norberg et al., 2012). Furthermore, current-driven dispersal patterns of coral larvae can potentially increase or decrease local bleaching thresholds (Kleypas et al. 2016). In our study, the cold configuration, which selected the coldest 30% of the network as MPAs, disproportionately protected destinations of warm-adapted larvae (i.e., preadapted to rising temperatures) but not sources. These sites benefitted from adaptive gene flow as long as sources of warm-adapted larvae maintained coral cover. Notably, the increase in cold-adapted larvae in the network led to a negative spillover effect in non-MPA sites, slowing adaptive evolution at those reefs. Conversely, the hot configuration, which selected the hottest 30% of the network as MPAs, disproportionately protected sources of warm-adapted larvae, but not destinations. These sites continued to benefit the rest of the network by providing adaptive gene flow, but did not substantively receive this benefit themselves, because few incoming larvae were preadapted to local temperatures.

Our findings suggest that, by protecting sites with particular temperature and connectivity characteristics, marine spatial planning may alter eco-evolutionary processes to enhance or inhibit the adaptive capacity of a reef network. Future work could test the likelihood and magnitude of these impacts through a combination of approaches, including metapopulation genetics, larval transplant or common garden experiments, long-term studies on the efficacy of local interventions to support coral health, and more realistic simulations enabled by advances in computation and data availability.

### Empirical gap analysis

We next assessed the extent to which existing MPAs represented important characteristics for climate change adaptation across three major coral reef regions. We found that current MPA networks are not fully representative of the thermal and connectivity characteristics of their broader regions. Rather, they have highly skewed temperature profiles. Like the hot configuration we simulated, temperature was higher in MPA sites and pr05, which measured warm connection strength, was lower than non-MPA sites in the Caribbean. Consequently, MPAs in the Caribbean likely do not receive as many warm-adapted larvae as non-MPAs. The opposite is true of the Southwest Pacific and Coral Triangle, which more closely approximated the cold MPA configuration: these MPA networks appear to receive more warm-adapted larvae than non-MPA sites.

The temperature profiles of existing MPAs in the Coral Triangle and Southwest Pacific are likely driven by the large number of MPAs in Australia’s Great Barrier Reef, which is located on the relatively cooler southern edge of both regions. Conversely, many of the existing MPAs in the Caribbean are located within the Meso-American Barrier Reef, which occupies relatively warm waters. Based on results from the eco-evolutionary simulations, the skewed temperature distributions and spatial clustering of protected habitats could imply that our focal MPA networks are not optimally configured to harness the adaptive potential of the full region. Namely, the existing Caribbean MPA network is poised to protect sources of warm-adapted larvae, but not destinations. The opposite is true of the Southwest Pacific and Coral Triangle. Eco-evolutionary theory suggests that regional coordination to facilitate the protection of diverse reef habitats may help to maintain the genetic diversity that fuels adaptive gene flow (Colton et al. 2022).

### Model limitations

Our model included several simplifying assumptions. We assumed that all reefs were composed of two spawning coral types and a macroalgae, ignoring potential impacts of more complex life histories and community structure, diversity, and local stressors. Notably, we did not include brooding species that are characterized by lower reproductive output and local-scale dispersal (Szmant 1986). Because brooders are self-seeding at the scale of our study, we expect that simulated reef systems dominated by these corals would result in a demographic boost that is similar across MPA configurations and would not result in the network-scale eco-evolutionary impacts that differentiate configurations in the spawning coral case. These dynamics would be most relevant for Caribbean reefs, where brooders compose approximately 57% of coral species. In contrast, brooders are relatively rare in the Indo-Pacific, composing approximately 5% of all species (Roff 2021). In general, including a greater number of functional types may further stabilize coral cover during the period of rapid temperature increase (Peterson et al. 1998; Winfree & Kremen 2009) because of processes related to the species diversity-stability hypothesis (Loreau et al. 2021). Life history parameterization of the model, in particular growth and death rates, led to hindcast coral cover within empirically observed ranges. Thus, while we expect our comparative MPA configuration results to hold across broad ranges of parameter values, uncertainties in rates due to a lack of relevant data means the years at which we predicted decline and recovery are only relative.

Because we simulated coral cover across all available coral reef habitat, MPAs in our scenarios may have been implemented in areas that would be considered too degraded for protection. Our interpretation of a protected area was also simplified, in that an MPA was represented as a site with higher macroalgal mortality. Therefore, MPAs in our model could represent both a reduction in fishing of herbivores and/or water quality improvement. Globally, MPAs differ in their objectives, regulations (Horta e Costa et al., 2016) and effectiveness (Rife et al. 2013). Protected areas that are designed to maximize fisheries biomass (Cabral et al. 2019) or reduce bycatch (Hastings et al. 2017), for example, may affect coral eco-evolutionary dynamics in counterintuitive ways that are not captured in the current model. Our model assumed that all MPA’s were perfectly implemented, ignoring heterogeneities in enforcement and governance; incorporating this variation in our model would likely decrease the effects of MPAs on the network since those reefs would be more similar to unprotected reefs. We also focused exclusively on the eco-evolutionary dynamics surrounding coral temperature tolerance and rising temperatures, and did not consider the impacts of other global and local stressors, such as sedimentation and ocean acidification.

Finally, we note that our model was implemented on large spatial scales, where reef size ranged from 5 to >100 km^2^, depending on the region. This scale dictated larval connectivity patterns, temperature distributions, and sizes of MPAs across the seascape. We made the simplifying assumption that each reef experiences a single mean temperature, with a range of phenotypic responses underlying the genetic variation within each reef. This assumption is based on two premises: 1) there exists a range of phenotypic responses within a reef site and 2) subpopulations within a reef site are more connected to each other than to subpopulations at other sites, leading to a collective population response to the site’s overall thermal environment. However, the range of phenotypic responses within a reef site in our model can also represent a range of corals adapted to local microclimates. An assessment of the impacts of intra-reef heterogeneity, including microclimates (Bay & Palumbi 2014) and smaller scales of dispersal, is beyond the scope of this paper. Because intra-reef heterogeneity may greatly affect local adaptation and the presence of adaptive alleles, this area of research is an exciting avenue for future modeling and field studies. We may be able to simulate the variation in thermal microclimate availability among sites in our model by adjusting genetic variance among sites. Such work would benefit from data on the distribution of microclimates across reefs, which currently is not available.

### Relevance to management and conclusions

While our study provides insights into general principles of MPA network design for coral persistence under climate change, it is important to note that real-world implementation would require consideration of additional local factors. These may include local residents’ desires and political will, socioeconomic considerations, existing management structures, specific local stressors, and finer-scale habitat heterogeneity (Weeks et al. 2010; Gill et al. 2017; Christie et al. 2017). These findings contribute to the toolkit for strategic planning and may be considered alongside detailed local knowledge as managers and policymakers design or expand MPA networks (Ban et al. 2011).

We used projected temperatures from the RCP 4.5 emissions scenario, which assumes that sufficient actions are taken to limit warming to between 2 and 3°C globally. A previous study showed that a more severe emissions scenario, RCP 8.5, led to coral cover declines across the three regions (McManus et al., 2021b). Here, even the RCP 4.5 scenario led to initial declines, indicating that reefs will “run the climate gauntlet” before temperatures stabilize (Hughes et al. 2017). As such, we observed coral recovery because of the reduced emissions scenario, underscoring that MPAs on their own are not sufficient to prevent large-scale coral declines.

In the RCP 4.5 simulations presented here, the differences in coral cover within and outside of MPAs among network configurations have significant implications for coral conservation and marine conservation. The goals of conservation practitioners and decision-makers can vary widely in both the types of benefits (i.e., ecosystem services or nature’s contributions to people) they seek to maintain or augment, as well as the focal spatial and temporal scales (O’Leary et al. 2016; Rilov et al. 2019; Wilson et al. 2020). We found that the representative configuration was best for spreading coral cover benefits across the seascape, both spatially and categorically, inside and outside of MPAs. Managers and conservation planners may be able to implement this approach by using temperature and connectivity information to identify underrepresented thermal and connectivity characteristics (e.g., cooler, warmer, source, and destinations sites), and prioritizing those sites for conservation going forward. Managers focused only on local benefits within MPAs and short-term benefits might choose the cold configuration, but this approach favors maladaptation and drives slower recovery after the climate stabilizes at the network scale. The current momentum to protect 30% of global oceans by 2030 (e.g., Chandrasekhar et al. 2022), combined with programs that echo this goal on national and subnational scales (e.g., Newsom 2020; The White House 2021) provide an opportunity for expanding MPA networks. In light of our results, decision-makers tasked with planning these expanded networks can consider the potential multi-scalar eco-evolutionary impacts and trade-offs of regional MPA network configurations. While we highlight important differences among MPA network configurations, we note that implementing protection at a given site—regardless of MPA network type—contributed to local coral persistence. Additionally, higher proportions of MPAs led to higher coral cover across the seascape. Together, these results highlight the importance of maintaining suitable local environmental conditions (Fabricius 2005; Wooldridge & Done 2009; De’ath et al. 2012; Suchley & Alvarez-Filip 2017; National Academies of Sciences, Engineering, and Medicine 2019; Donovan et al. 2021; Gove et al. 2023) and suggest that expanding conservation or management efforts that enhance coral growth will benefit coral cover at both local and seascape scales.

Our work shows that the distribution, extent, and effectiveness of local interventions have the potential to affect the distribution of coral cover beyond what would be expected from local benefits alone, due to the potentially wide-reaching effects of larval dispersal and gene flow. However, the rate and duration of ocean warming and local stressors are the primary drivers of the decline of coral reefs (Bay et al. 2017; McManus et al. 2020, 2021b; Matz et al. 2020; Logan et al. 2021). Therefore, limiting the degree of ocean warming via reductions in greenhouse gas emissions and reducing local stressors at a greater number of sites (by expanding MPAs, improving water quality, or other means of intervention) are important for coral conservation, independent of any particular MPA configuration explored in this paper. Our research further emphasizes that the coming decades are a critical time for the conservation of coral reefs. This work suggests that the future distribution of coral reefs likely rests on a combination of regionally-coordinated local interventions *and* emissions reductions.

## Acknowledgements

We gratefully acknowledge funding from the Gordon and Betty Moore Foundation and The Nature Conservancy.

Additional supporting information may be found in the online version of the article at the publisher’s website.

