## Supplemental Information for "Marine spatial planning to enhance coral adaptive potential"

### Supporting Information

#### Appendix S1: Eco-evolutionary model details

On each reef, we track the change in coral cover of each type and their mean optimal growth temperatures. The cover of coral type $i$ on reef patch $a$, $N_{i,a}$, responds to population growth ($g_{i,a}$), larval immigration, and genetic load (Eq. 1). The larval recruitment rate $(l_{i,a})$ is scaled by the remaining free space on the patch, $F_a$, such that recruitment is zero when there is no space for settlement ($F_a$=0). The additive genetic variance ($V$) sets the rate of evolution. Higher $V$ allows a faster evolutionary response, but also increases the genetic load, which is a cost for the presence of individuals that are suboptimally adapted (Kirkpatrick & Barton 1997). Finally, $z_{i,a}$ is the mean trait value of each type, defined here as the optimal temperature for growth. Together, these processes are modeled as:

$$\frac{\mathrm{d}N_{i,a}}{\mathrm{d}t} = g_{i,a}N_{i,a} + l_{i,a}F_a + \frac{1}{2}V\frac{\partial^2 g_{i,a}}{\partial z^2}\bigg|_{z=z_{i,a}} N_{i,a} \tag{1}$$

[change in coral cover] = [population growth] + [larval immigration] + [genetic load]

We also track the change in the mean trait value — the mean thermal optimum — at each patch, $z_{i,a}$. This trait responds to gene flow and stabilizing selection (Eq. 2). The trait value of larvae arriving via dispersal to each reef ($y_{i,a}$) is scaled by the proportion of coral cover represented by newly-settled larvae, $l_{i,a}/N_{i,a}$, and by available free space $F_a$. Stabilizing selection, $q_i V \frac{\partial g_{i,a}}{\partial z}$, drives the mean trait value towards the local temperature; higher $V$ increases the rate of this evolutionary response (Lande 1976). When coral cover is exceedingly low, $q_i$ reduces the rate of evolution because very small populations tend to lose genetic variance (Williams et al. 2018). The equation governing the mean trait value for each reef is written as:

$$\frac{\mathrm{d}z_{i,a}}{\mathrm{d}t} = \left(y_{i,a} - z_{i,a}\right)\left(\frac{l_{i,a}}{N_{i,a}}\right)F_a + q_i V\frac{\partial g_{i,a}}{\partial z}\bigg|_{z=z_{i,a}} \tag{2}$$

[change in optimal growth temperature] = [gene flow] + [stabilizing selection]

Local population dynamics are governed by the overall fitness of the population, $g_i$ (Eq. 3), where $r_i$ is the intrinsic growth rate, $m_i$ is the mortality rate, and $\alpha_{ij}$ is the competitive effect of type $j$ on type $i$; the matrix $\boldsymbol{\alpha}$ comprised these competitive interactions.

$$g_{i,a} = r_{i,a}\left(1 - \sum_j \alpha_{ij} N_{j,a}\right) - m_{i,a} \tag{3}$$

Intrinsic growth (Eq. 4) and mortality (Eq. 5) are modeled as Gaussian and exponential functions, respectively, of the difference between local temperature, $T_a$, and mean optimal growth temperature, $z_{i,a}$. These rates are also dependent on the width of thermal tolerance of each type, $w_{i,}$ and their growth scaling factor, $r_{0,i}$. Because many thermal performance curves are left-skewed (Deutsch et al. 2008), we increase mortality when the local temperature is hotter than the mean optimal growth temperature, $T_a > z_{i,a}$.

$$r_{i,a} = \frac{r_{0,i}}{\sqrt{2\pi w_i^2}} exp\left(\frac{-(T_a - z_{i,a})^2}{w_i^2}\right) \tag{4}$$

$$m_{i,a} = \begin{cases} 0, & T_a \leq z_{i,a} \\ 1 - exp\left(\frac{-(T_a - z_{i,a})^2}{w_i^2}\right), & T_a > z_{i,a} \end{cases} \tag{5}$$

Following (McManus et al. 2021), we model spatially explicit larval dispersal among reef patches. The rate of larval input for each reef *a* is dependent on the coral cover at each source reef, the effective fecundity rate ($\beta$), and the connectivity matrix $\mathbf{D}$, where element $D_{ab}$ is the probability of dispersal from patch $b$ to patch $a$. Larval input also depends on the area of focal reef $a$, and the areas of each source reef $b$; these values are denoted as $A_a$ and $A_b$, respectively. The amount of free space available ($F_a$) is calculated from the proportional cover of each type.

$$l_{i,a} = \frac{\beta}{A_a} \sum_b A_b \mathrm{D}_{ab} N_{i,b} \tag{6}$$

$$F_a = 1 - \sum_i N_{i,a} \tag{7}$$

In addition to demographic effects, larval immigration affects the mean optimal growth temperature through gene flow (first term on right-hand side of Eq. 2). On each reef, the mean incoming trait value is calculated as

$$y_{i,a} = \frac{\frac{\beta}{A_a} \sum_b A_b \mathrm{D}_{ab} N_{i,b} z_{i,b}}{l_{i,a}} = \frac{\sum_b A_b \mathrm{D}_{ab} N_{i,b} z_{i,b}}{\sum_b A_b \mathrm{D}_{ab} N_{i,b}} \quad (8)$$

At very low coral cover ($N_{min} = 10^{-6}$), $q_i$ reduces the effects of selection (Norberg et al. 2012).

$$q_i = \max\left(0, 1 - \frac{N_{min}}{\max(N_{min}, 2N_i)}\right) \quad (9)$$

Macroalgae, denoted as type *M*, are included in the framework to serve as a competitor for corals and to facilitate the implementation of MPAs (Eqs. 10-12). As such, macroalgal growth, $r_{M,0}$ and mortality, $m_{M,a}$, are insensitive to temperature and set as constants. Macroalgae also do not experience evolutionary effects nor dispersal among patches. However, macroalgae mortality is higher at MPA reefs relative to those that are not designated as MPAs. For macroalgae, only the first term of Eq. 1 is relevant while Eq. 2 is not applicable. The population dynamics of macroalgae are then tracked as:

$$\frac{\mathrm{d}N_{M,a}}{\mathrm{d}t} = g_{M,a} N_{M,a} \quad (10)$$

$$g_{M,a} = r_{M,0}\left(1 - \sum_j \alpha_{ij} N_{j,a}\right) - m_{M,a} \quad (11)$$

$$m_{M,a} = \begin{cases} 0.3, & \text{outside MPA} \\ 0.5, & \text{inside MPA} \end{cases} \quad (12)$$

#### Supporting tables and figures

**Appendix S2.** Parameter definitions and values used in simulations for the two coral types *i*.

| Parameter | Definition | | Fast coral | Slow coral | Macroalgae |
|---|---|---|---|---|---|
| $r_{0,i}$ | Scaling factor for growth rate; Note that maximum growth rate is $\frac{r_{0,i}}{\sqrt{2\pi w^2}}$ | | 1.5 | 1.5 | N/A |
| $w_i$ | Thermal tolerance breadth | | 1.0 | 3.0 | N/A |
| $\beta$ | Effective fecundity | | 0.5 | 0.5 | N/A |
| $V$ | Additive genetic variance | | 0.01 | 0.01 | N/A |
| $r_M$ | Macroalgal growth rate | | N/A | N/A | 1.0 |
| $m_M$ | Macroalgal mortality rate | | N/A | N/A | outside MPA: 0.3<br>inside MPA: 0.5 |
| $\alpha$ | Competition matrix | Fast coral | 5.77 | 0.9 | 1.0 |
| | | Slow coral | 0.9 | 5.77 | 1.0 |
| | | Macroalgae | 1.0 | 1.0 | 1.0 |

**Appendix S3**. Methods and parameters used to create the connectivity matrices.

| **Item Description** | **Caribbean** | **Southwest Pacific** | **Coral Triangle** |
|---|---|---|---|
| Publication | Schill et al. 2015 | Treml et al. 2008 | Thompson et al. 2018 |
| Number of Reef Units | 423 | 583 | 2083 |
| Reef location and area determination | Millenium Coral Reef Mapping Project for reef locations and areas, reviewed by in-country reef experts | Digital Chart of the World Server plus Spalding et al. 2001 & Oliver et al. 2004 | Global Distribution of Coral Reefs (UNEP-WCMC), which merges data from the Millennium Coral Reef Mapping Project and the World Atlas of Coral Reefs |
| Ocean Circulation Model | NOAA Real-Time Ocean Forecast System [RTOFS] database (8 x 8 km resolution) (Mehra & Rivin 2010) | NOAA Environmental Modeling Center's ocean analysis system (25 x 25 km resolution) (Ji et al. 1995) | Regional Ocean Modelling System developed for the Coral Triangle region [CT-ROMS] (~5 x 5 km resolution) (Castruccio et al. 2013) |
| Maximum Pelagic Larval Duration | 30 days | 60 days | 30 days |
| Larval Mortality | 20% per day | 4% per day | none |
| Spawning events | Two per year from 2008-2011, starting on the last dates of the quarter moon (August - September) | One mass spawning season - September through November 2001 | Biannual mass spawning events lasting 5 days (spring and fall) |
| Pre-competency Period | Gamma cumulative function allowing all larvae to reach full competency in 3 days | none | Beginning at 3 days with full competency at 10 days |

| | | | |
|---|---|---|---|
| Settlement behavior | After reaching competency, larvae over coral habitat settled at a rate of 75% per day. | If the density of larvae at a downstream reef site exceeded 1 per cell, a connection was made between the two sites | none |
| Local density & fecundity | Amount of larvae released proportional to reef area | 10,000 larvae per $km^2$ | 25 particles from each of the oceanic grid cells within a reef site, for a maximum of 8000 particles released from each site. |

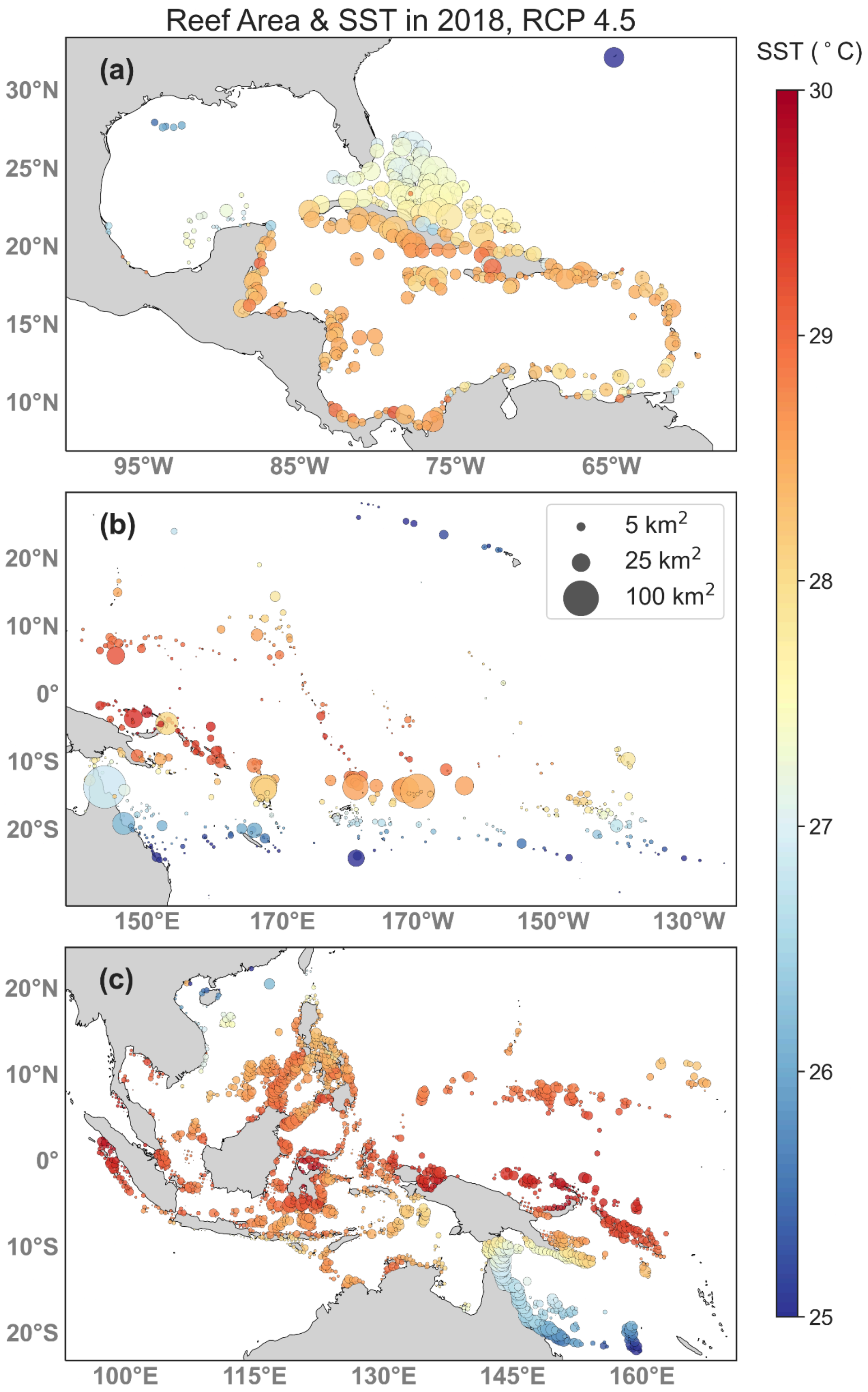
Reef Area & SST in 2018, RCP 4.5
SST (°C)
30
29
28
27
26
25
(a)
30°N
25°N
20°N
15°N
10°N
95°W
85°W
75°W
65°W
(b)
5 km$^2$
25 km$^2$
100 km$^2$
20°N
10°N
0°
10°S
20°S
150°E
170°E
170°W
150°W
130°W
(c)
20°N
10°N
0°
10°S
20°S
100°E
115°E
130°E
145°E
160°E

**Appendix S4.** Maps of reef units scaled by area, and colored by temperature in the Caribbean (a), Southwest Pacific (b), and Coral Triangle (c).

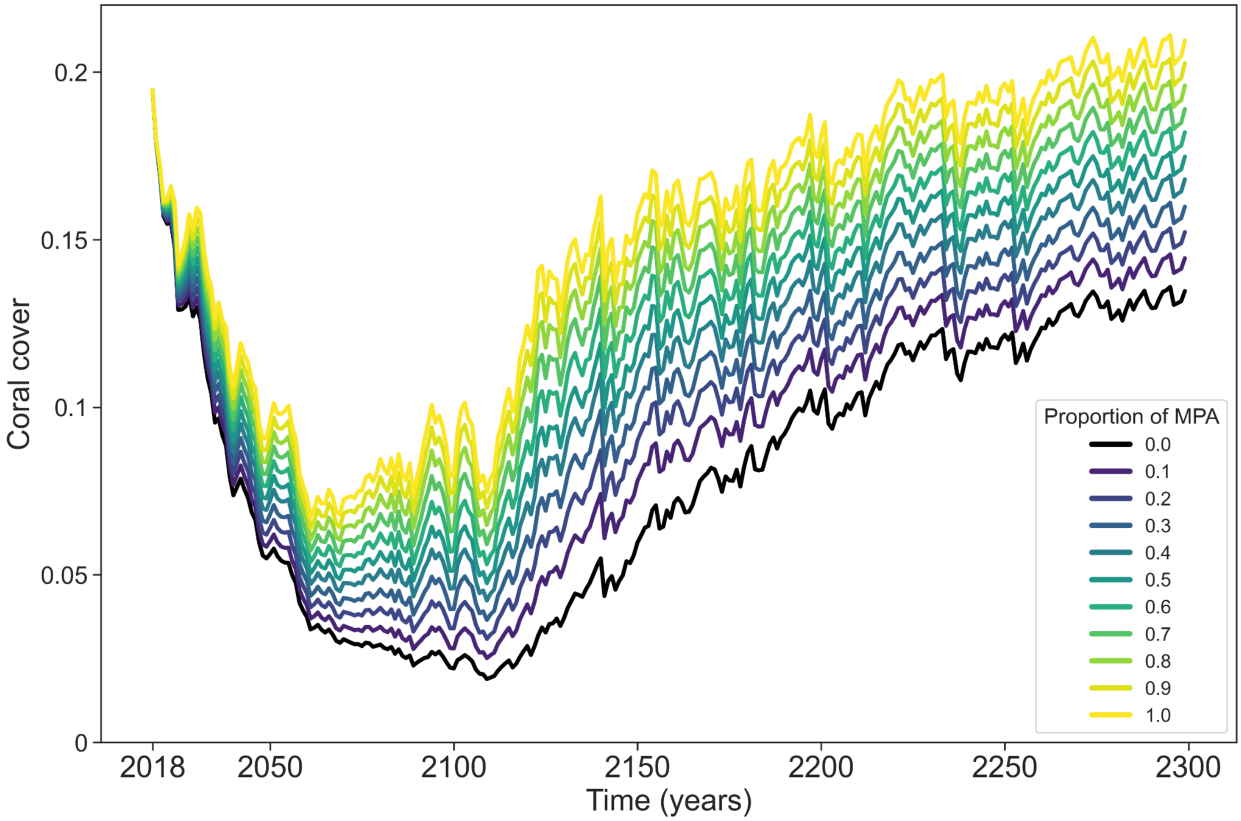


**Appendix S5.** Mean coral cover through time in the Caribbean with the representative MPA configuration. Line colors represent different levels of MPA coverage. In the main text, 30% of sites are included in the MPA network.

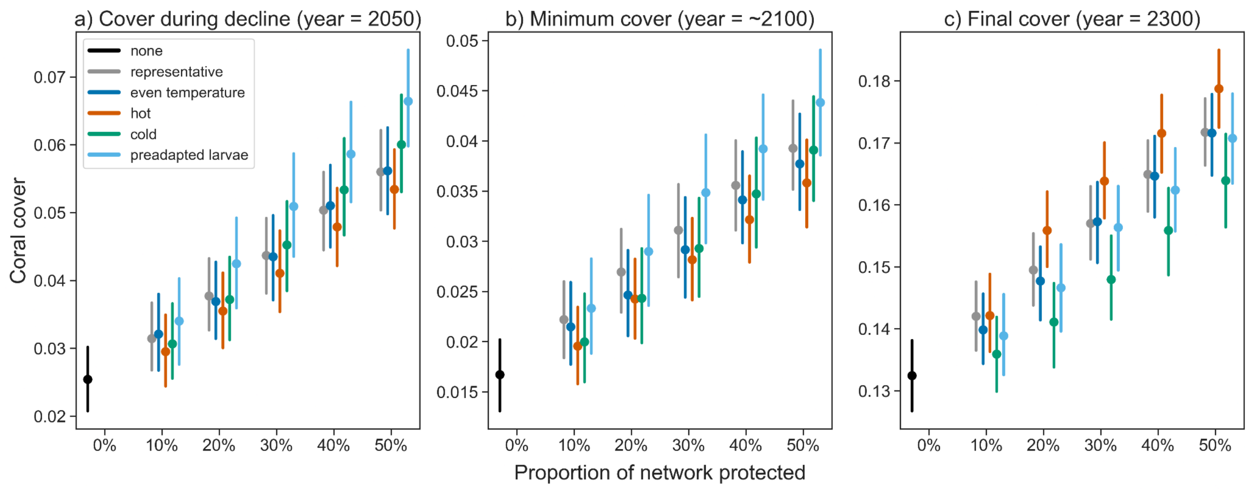


**Appendix S6.** Mean coral cover in the Caribbean under different MPA network configurations from 0 (none) to 50% of reefs designated as MPAs. Panel a shows cover during decline (year = 2050), b shows minimum coral cover, and c shows cover at the end of the simulation (year = 2300).

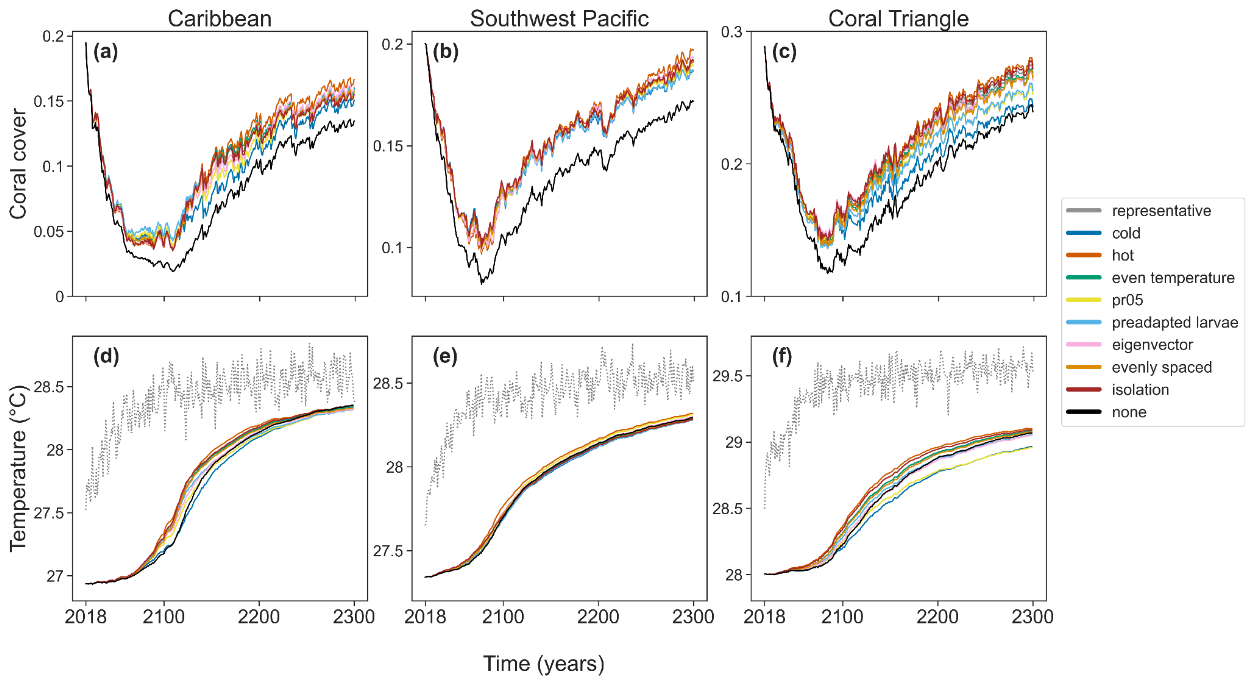


**Appendix S7.** Trajectories of mean total coral cover and mean trait values for all MPA network configurations. Gray dotted lines in d-f indicate the projected mean network-wide sea surface temperature time series.

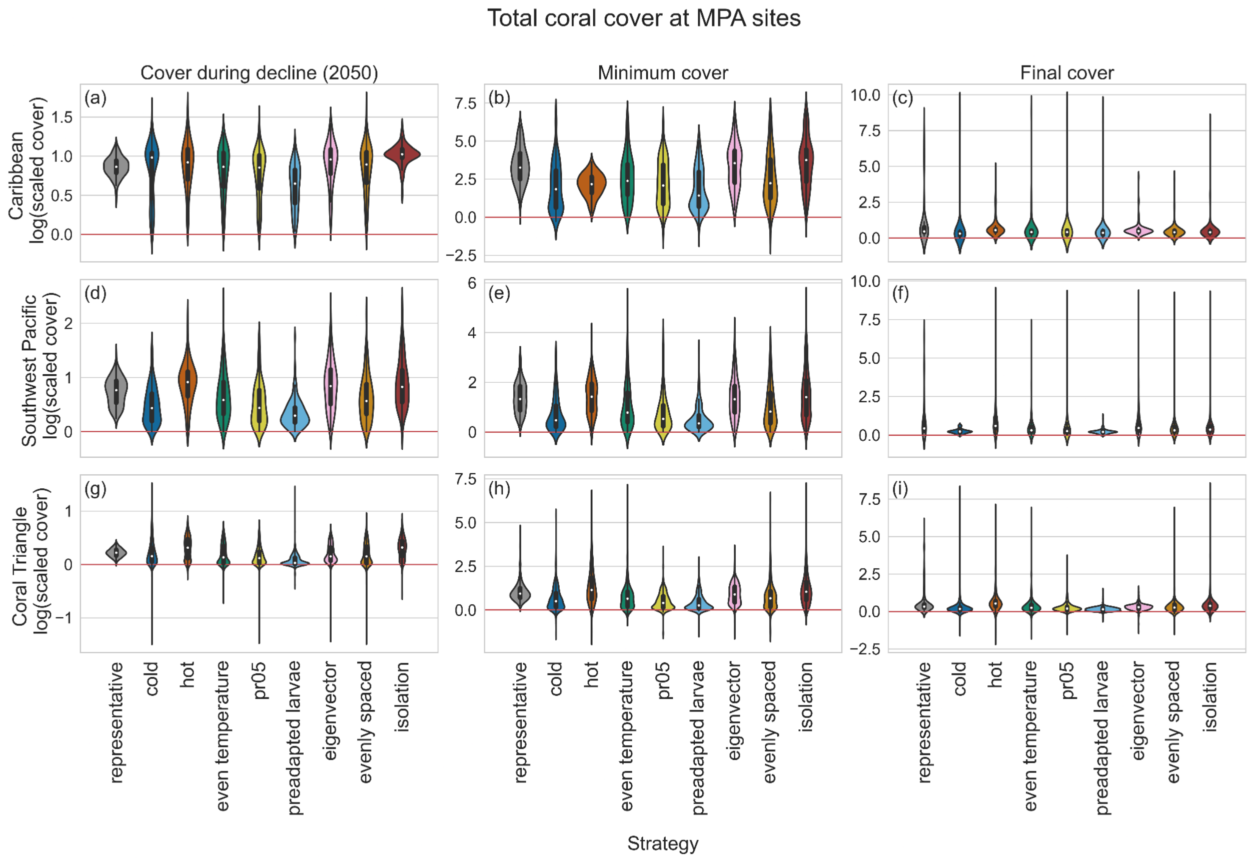


**Appendix S8.** Coral cover at MPA sites in the Caribbean (A-C), Southwest Pacific (D-F) and Coral Triangle (G-I) under each configuration scaled by cover when there were no MPAs. Results above the red horizontal line indicate strategies where coral cover was higher than in the simulations with no MPAs while results below the line indicate the opposite. Means are marked as white dots.

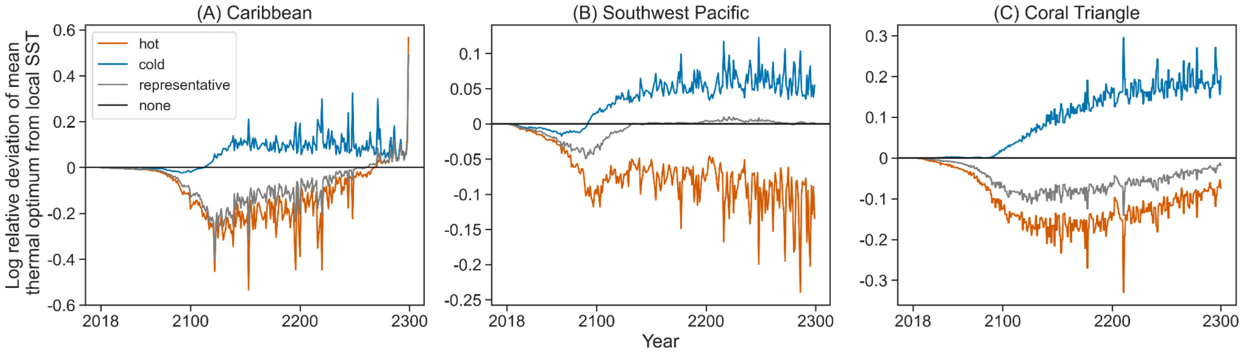


**Appendix S9.** The absolute deviation of the mean trait value from local SST under hot (orange), cold (blue), representative (gray) MPA configurations, relative to the deviation under the scenario without MPAs. Results above the black horizontal line indicate that the deviation is greater than without MPAs, and results below the line indicate that the deviation is smaller than without MPAs.

**Appendix S10. Caribbean coral cover at multiple time points.** Cover at decline is taken in the year 2050.

| MPA strategy | cover at decline | minimum cover | final cover | inside cover at decline | inside minimum cover | inside final cover | outside cover at decline | outside minimum cover | outside final cover |
|---|---|---|---|---|---|---|---|---|---|
| representative | 0.068 | 0.036 | 0.16 | 0.096 | 0.062 | 0.203 | 0.055 | 0.024 | 0.141 |
| cold | 0.069 | 0.035 | 0.151 | 0.134 | 0.087 | 0.235 | 0.042 | 0.008 | 0.115 |
| hot | 0.065 | 0.036 | 0.167 | 0.076 | 0.047 | 0.19 | 0.06 | 0.03 | 0.157 |
| even temperature | 0.067 | 0.036 | 0.16 | 0.102 | 0.068 | 0.206 | 0.053 | 0.022 | 0.14 |
| pr05 | 0.07 | 0.038 | 0.157 | 0.129 | 0.088 | 0.225 | 0.045 | 0.015 | 0.128 |
| preadapted larvae | 0.071 | 0.043 | 0.159 | 0.158 | 0.11 | 0.247 | 0.034 | 0.009 | 0.122 |
| eigenvector centrality | 0.065 | 0.037 | 0.161 | 0.074 | 0.048 | 0.191 | 0.061 | 0.027 | 0.149 |
| evenly spaced | 0.067 | 0.035 | 0.157 | 0.096 | 0.065 | 0.194 | 0.055 | 0.022 | 0.142 |
| isolation | 0.068 | 0.035 | 0.155 | 0.074 | 0.041 | 0.186 | 0.065 | 0.026 | 0.142 |
| none | 0.054 | 0.019 | 0.135 | 0.054 | 0.019 | 0.135 | 0.054 | 0.019 | 0.135 |

**Appendix S11. Southwest Pacific coral cover at multiple time points.** Cover at decline is taken in the year 2050.

| MPA strategy | cover at decline | minimum cover | final cover | inside cover at decline | inside minimum cover | inside final cover | outside cover at decline | outside minimum cover | outside final cover |
|---|---|---|---|---|---|---|---|---|---|
| representative | 0.115 | 0.099 | 0.192 | 0.145 | 0.134 | 0.231 | 0.102 | 0.084 | 0.175 |
| cold | 0.115 | 0.098 | 0.187 | 0.19 | 0.183 | 0.272 | 0.083 | 0.062 | 0.15 |
| hot | 0.115 | 0.097 | 0.197 | 0.099 | 0.067 | 0.179 | 0.121 | 0.106 | 0.204 |
| even temperature | 0.115 | 0.098 | 0.192 | 0.149 | 0.135 | 0.232 | 0.101 | 0.083 | 0.175 |
| pr05 | 0.114 | 0.098 | 0.189 | 0.178 | 0.169 | 0.252 | 0.087 | 0.068 | 0.163 |
| preadapted larvae | 0.116 | 0.1 | 0.187 | 0.228 | 0.213 | 0.285 | 0.068 | 0.052 | 0.145 |
| eigenvector centrality | 0.115 | 0.098 | 0.193 | 0.122 | 0.096 | 0.199 | 0.113 | 0.096 | 0.191 |
| evenly spaced | 0.116 | 0.099 | 0.191 | 0.153 | 0.14 | 0.237 | 0.099 | 0.082 | 0.171 |
| isolation | 0.115 | 0.1 | 0.192 | 0.107 | 0.1 | 0.211 | 0.119 | 0.099 | 0.184 |
| none | 0.1 | 0.082 | 0.172 | 0.1 | 0.082 | 0.172 | 0.1 | 0.082 | 0.172 |

**Appendix S12. Coral Triangle coral cover at multiple time points.** Cover at decline is taken in the year 2050.

| MPA strategy | cover at decline | minimum cover | final cover | inside cover at decline | inside minimum cover | inside final cover | outside cover at decline | outside minimum cover | outside final cover |
|---|---|---|---|---|---|---|---|---|---|
| representative | 0.226 | 0.141 | 0.269 | 0.241 | 0.16 | 0.294 | 0.22 | 0.132 | 0.258 |
| cold | 0.227 | 0.138 | 0.243 | 0.279 | 0.211 | 0.33 | 0.205 | 0.098 | 0.206 |
| hot | 0.226 | 0.136 | 0.274 | 0.19 | 0.103 | 0.244 | 0.242 | 0.151 | 0.287 |
| even temperature | 0.225 | 0.139 | 0.266 | 0.251 | 0.175 | 0.305 | 0.214 | 0.124 | 0.249 |
| pr05 | 0.226 | 0.141 | 0.252 | 0.288 | 0.215 | 0.325 | 0.199 | 0.106 | 0.221 |
| preadapted larvae | 0.222 | 0.136 | 0.255 | 0.335 | 0.247 | 0.372 | 0.174 | 0.088 | 0.205 |
| eigenvector centrality | 0.227 | 0.144 | 0.263 | 0.263 | 0.181 | 0.324 | 0.211 | 0.125 | 0.237 |
| evenly spaced | 0.225 | 0.138 | 0.265 | 0.242 | 0.165 | 0.297 | 0.217 | 0.126 | 0.251 |
| isolation | 0.228 | 0.142 | 0.272 | 0.184 | 0.115 | 0.241 | 0.247 | 0.152 | 0.285 |
| none | 0.216 | 0.117 | 0.239 | 0.216 | 0.117 | 0.239 | 0.216 | 0.117 | 0.239 |

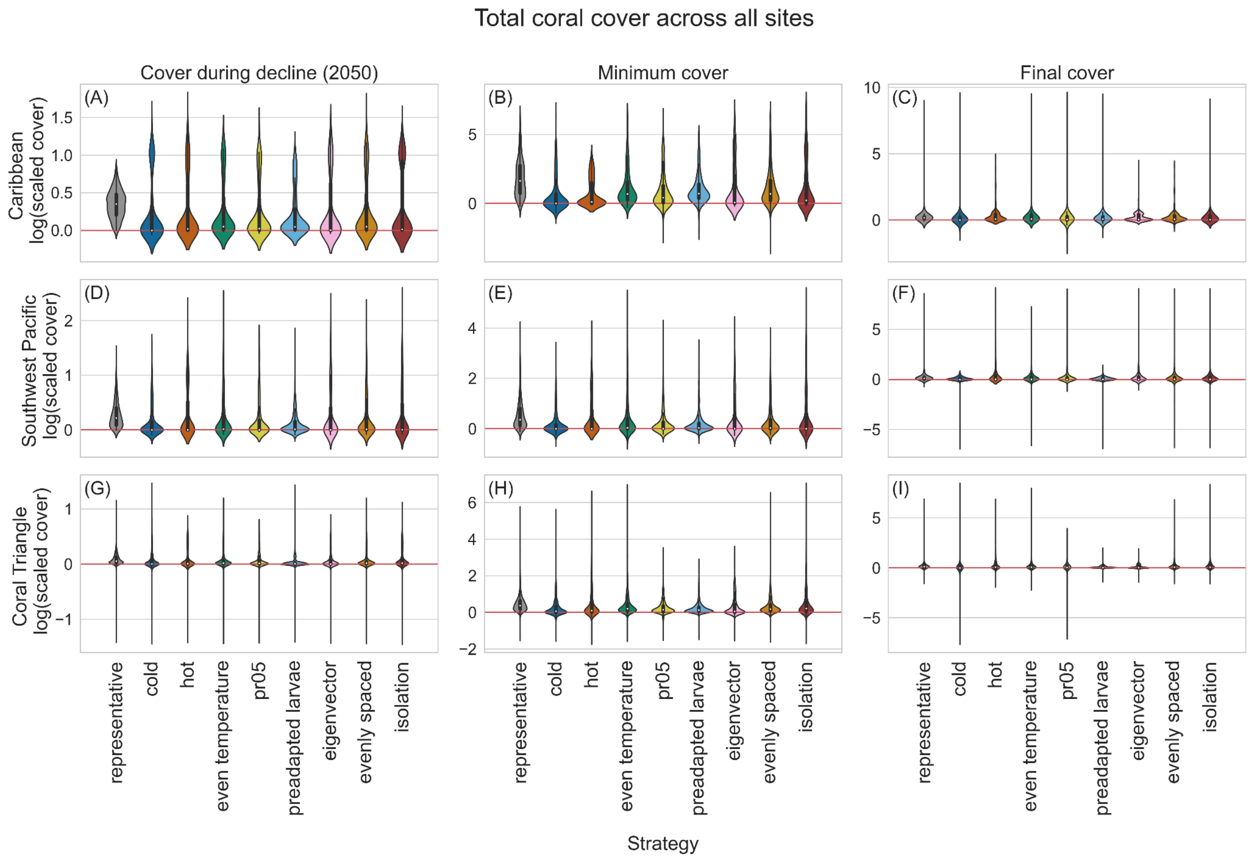


**Appendix S13.** Total coral cover for all sites (MPA + non-MPA sites) in the Caribbean (A-C), Southwest Pacific (D-F) and Coral Triangle (G-I) under each MPA strategy, scaled by cover when there were no MPAs. Log(relative coral cover) (y-axis) was calculated by dividing the cover at each site under a particular MPA scenario by the same site's cover when there were no MPAs and taking the log of that value. Results above the red horizontal line indicate strategies where coral cover was higher than in the simulations with no MPAs while results below the line indicate the opposite. Means are marked as white dots.

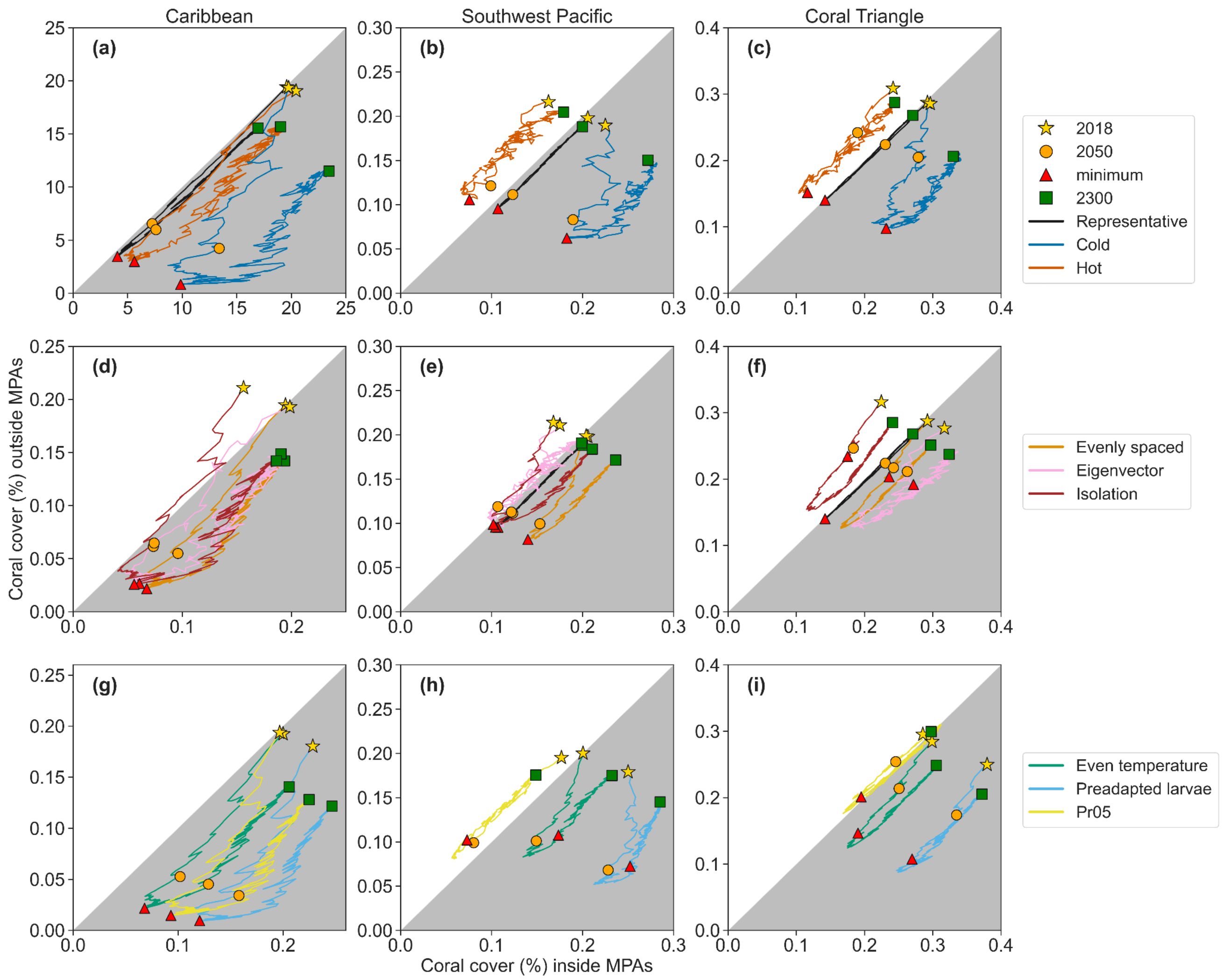


**Appendix S14.** Trajectories of mean total coral cover outside MPAs (y-axis) vs. inside MPAs (x-axis) for representative, cold, and hot configurations (top row); evenly spaced, eigenvector centrality and isolation configurations (middle row); and even temperature, preadapted larvae and pr05 configurations (bottom row). Trajectories in the white upper triangle indicate higher cover in sites not designated as MPAs as compared to MPA sites, while trajectories in the gray lower triangle reflect the opposite.

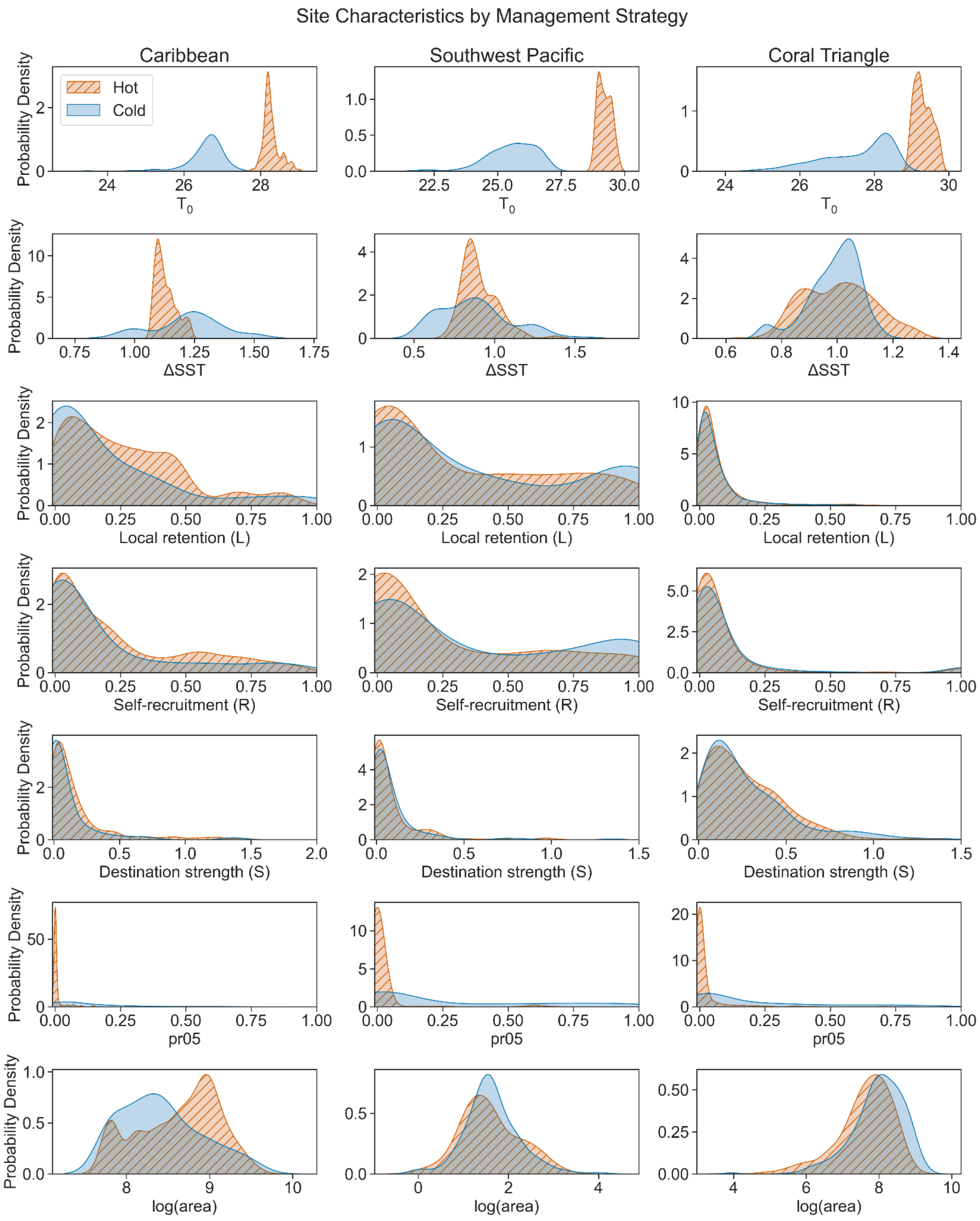
Site Characteristics by Management Strategy
Caribbean
Southwest Pacific
Coral Triangle
Hot
Cold
Probability Density
T0
ΔSST
Local retention (L)
Self-recruitment (R)
Destination strength (S)
pr05
log(area)

**Appendix S15.** Kernel density functions for hot and cold network configuration sites across all regions (columns) of the following site characteristics (rows, top to bottom): initial sea surface temperature ($T_0$), change in sea surface temperature ($\Delta T$), local retention ($L$), self-recruitment ($R$), destination strength ($S$), pr05 and log(area).

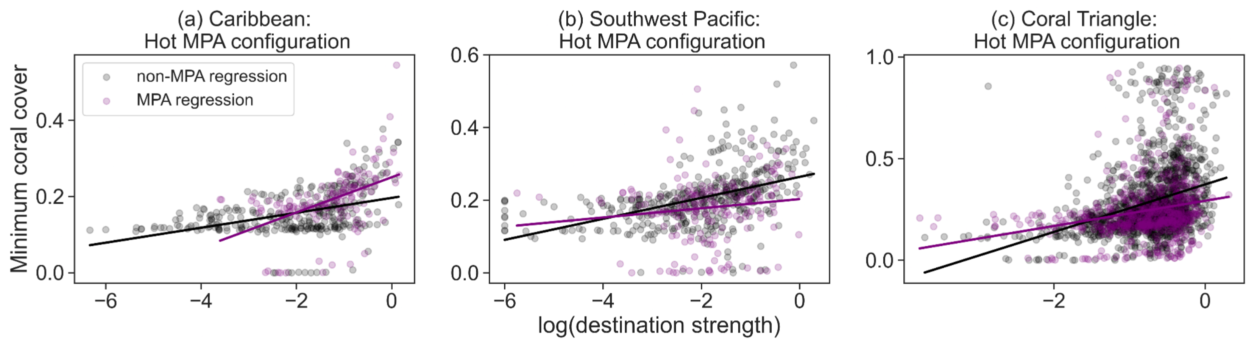


**Appendix S15.** Linear regression plots showing the relationship between log(destination strength) and minimum coral cover for MPA (purple dots) and non-MPA sites (black dots).

**Appendix S17**. Multiple regression results with minimum coral cover in the Caribbean under the representative MPA configuration (30%) as the response variable. Results show coefficient values +/- standard error for the three predictor variables: initial sea surface temperature, destination strength and MPA status. Results are shown for each representative iteration since each iteration selected a different subset of reefs. (P-value <0.05 = *, <0.01=**, <0.001 =***)

| | **Initial SST** | **Destination strength** | **MPA status** |
|---|---|---|---|
| **Random iteration** | Coefficient values ± standard error | | |
| 1 | -0.3521 ± 0.038*** | 0.4422 ± 0.038*** | 0.2934 ± 0.038*** |
| 2 | -0.3759 ± 0.039*** | 0.4458 ± 0.039*** | 0.2668 ± 0.039*** |
| 3 | -0.3781 ± 0.038*** | 0.4758 ± 0.038*** | 0.2591 ± 0.038*** |
| 4 | -0.3233 ± 0.039*** | 0.4402 ± 0.039*** | 0.3254 ± 0.039*** |
| 5 | -0.3786 ± 0.039*** | 0.4168 ± 0.039*** | 0.3165 ± 0.039*** |
| 6 | -0.3866 ± 0.039*** | 0.4271 ± 0.039*** | 0.2527 ± 0.039*** |
| 7 | -0.3520 ± 0.038*** | 0.4196 ± 0.038*** | 0.3418 ± 0.038*** |
| 8 | -0.3720 ± 0.040*** | 0.4444 ± 0.039*** | 0.3093 ± 0.040*** |
| 9 | -0.3721 ± 0.038*** | 0.4252 ± 0.038*** | 0.2943 ± 0.038*** |
| 10 | -0.3868 ± 0.038*** | 0.4358 ± 0.038*** | 0.3047 ± 0.038*** |

**Appendix S18**. Multiple regression results with minimum coral cover in the Southwest Pacific under the representative MPA configuration (30%) as the response variable. Results show coefficient values +/- standard error for the three predictor variables: initial sea surface temperature, destination strength and MPA status. Results are shown for each representative iteration since each iteration selected a different subset of reefs. (P-value <0.05 = *, <0.01=**, <0.001 =***)

| | **Initial SST** | **Destination strength** | **MPA status** |
|---|---|---|---|
| **Random iteration** | Coefficient values ± standard error | | |
| 1 | -0.3907 ± 0.035*** | 0.3355 ± 0.035*** | 0.2589 ± 0.034*** |
| 2 | -0.4091 ± 0.034*** | 0.3436 ± 0.034*** | 0.1935 ± 0.034*** |
| 3 | -0.4107 ± 0.035*** | 0.3497 ± 0.035*** | 0.1617 ± 0.035*** |
| 4 | -0.4056 ± 0.035*** | 0.3407 ± 0.035*** | 0.1862 ± 0.035*** |
| 5 | -0.3964 ± 0.035*** | 0.3253 ± 0.035*** | 0.2268 ± 0.035*** |
| 6 | -0.4085 ± 0.035*** | 0.3366 ± 0.035*** | 0.1995 ± 0.035*** |
| 7 | -0.4069 ± 0.035*** | 0.3348 ± 0.035*** | 0.1726 ± 0.035*** |
| 8 | -0.4106 ± 0.034*** | 0.3486 ± 0.034*** | 0.2406 ± 0.034*** |
| 9 | -0.4008 ± 0.035*** | 0.3316 ± 0.035*** | 0.2025 ± 0.035*** |
| 10 | -0.3946 ± 0.035*** | 0.3409 ± 0.035*** | 0.2130 ± 0.035*** |

**Appendix S19**. Multiple regression results with minimum coral cover in the Coral Triangle under the representative MPA configuration (30%) as the response variable. Results show coefficient values +/- standard error for the three predictor variables: initial sea surface temperature, destination strength and MPA status. Results are shown for each representative iteration since each iteration selected a different subset of reefs. (P-value <0.05 = *, <0.01=**, <0.001 =***)

| | **Initial SST** | **Destination strength** | **MPA status** |
|---|---|---|---|
| **Random iteration** | Coefficient values ± standard error | | |
| 1 | -0.3087 ± 0.020*** | 0.2735 ± 0.020*** | 0.1085 ± 0.020*** |
| 2 | -0.3021 ± 0.020*** | 0.2751 ± 0.020*** | 0.0974 ± 0.020*** |
| 3 | -0.3022 ± 0.020*** | 0.2725 ± 0.020*** | 0.0997 ± 0.020*** |
| 4 | -0.3093 ± 0.020*** | 0.2714 ± 0.020*** | 0.1254 ± 0.020*** |
| 5 | -0.2924 ± 0.020*** | 0.2893 ± 0.020*** | 0.0818 ± 0.020*** |
| 6 | -0.3072 ± 0.020*** | 0.2837 ± 0.020*** | 0.0892 ± 0.020*** |
| 7 | -0.2968 ± 0.020*** | 0.2783 ± 0.020*** | 0.0976 ± 0.020*** |
| 8 | -0.3059 ± 0.020*** | 0.2744 ± 0.020*** | 0.0760 ± 0.020*** |
| 9 | -0.3017 ± 0.020*** | 0.2747 ± 0.020*** | 0.0936 ± 0.020*** |
| 10 | -0.2928 ± 0.020*** | 0.2784 ± 0.020*** | 0.0839 ± 0.020*** |

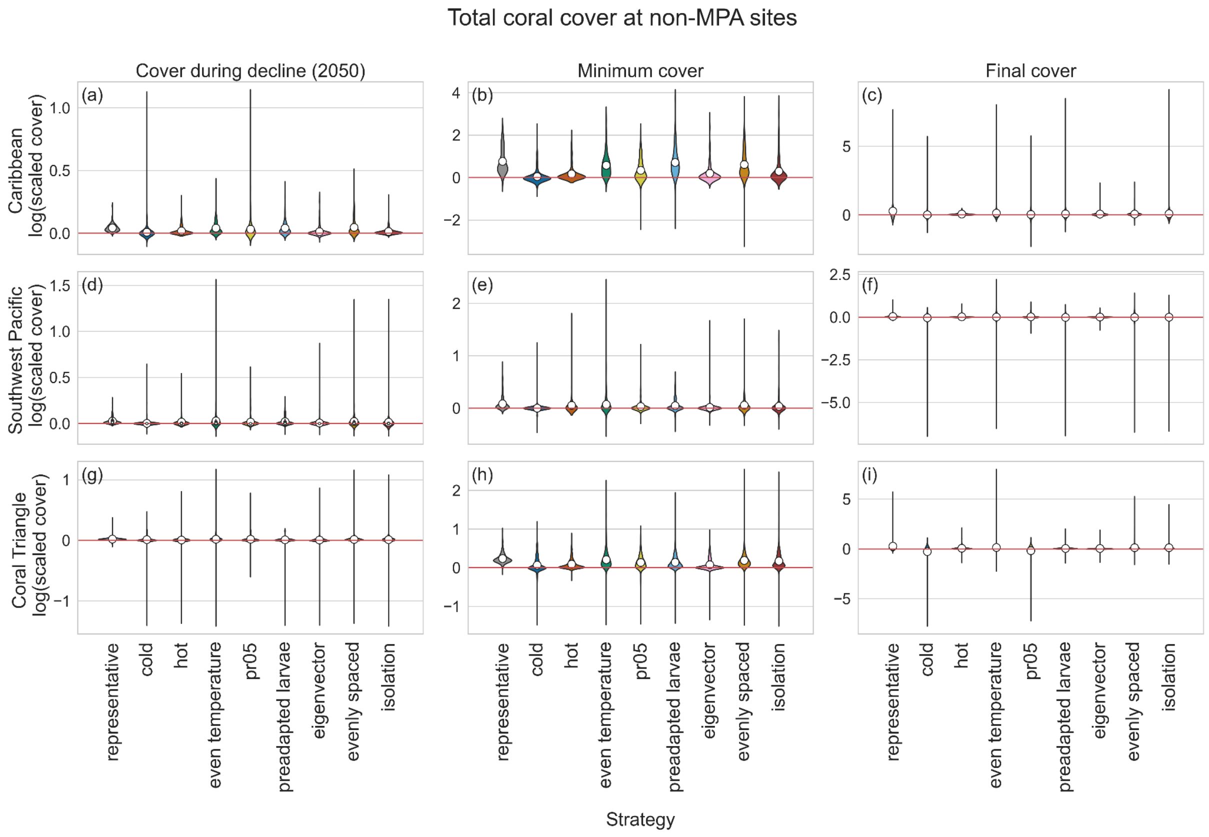
Total coral cover at non-MPA sites
Cover during decline (2050)
Minimum cover
Final cover
Caribbean log(scaled cover)
Southwest Pacific log(scaled cover)
Coral Triangle log(scaled cover)
(a)
(b)
(c)
(d)
(e)
(f)
(g)
(h)
(i)
representative
cold
hot
even temperature
pr05
preadapted larvae
eigenvector
evenly spaced
isolation
Strategy

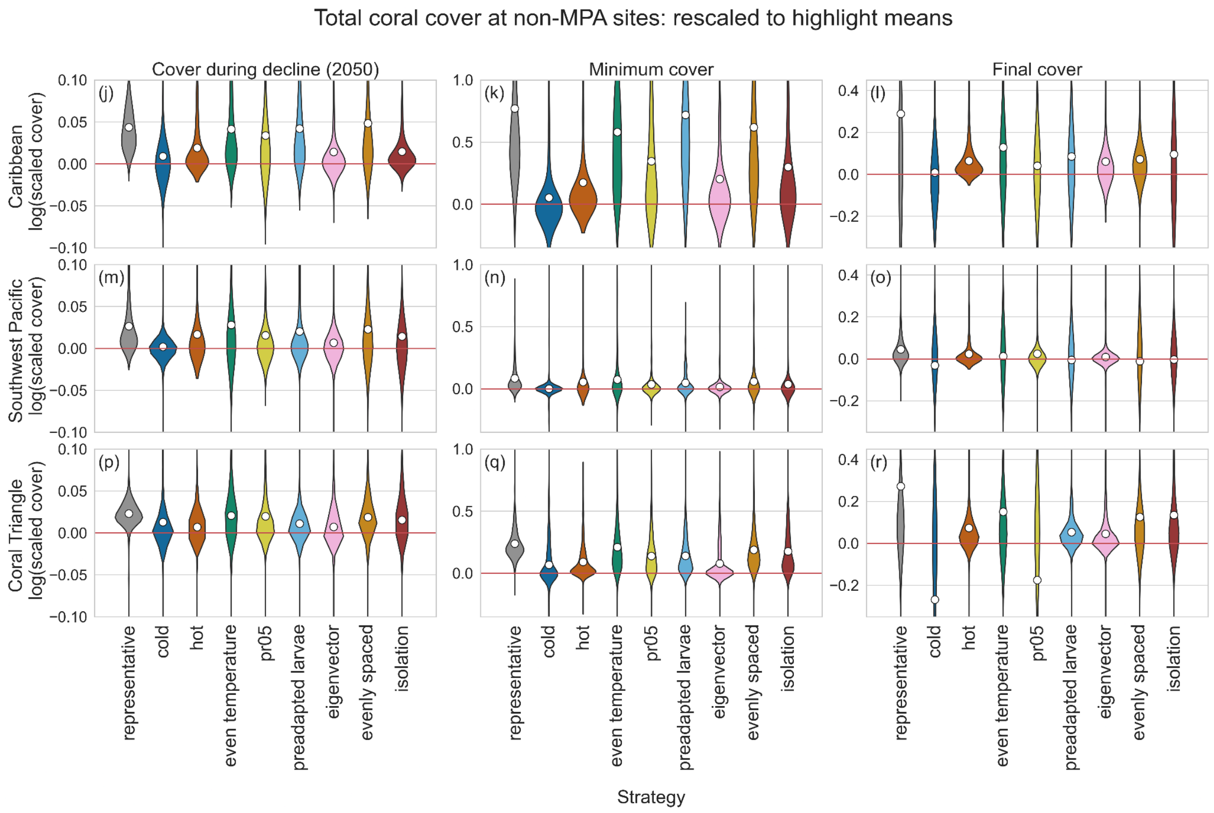
Total coral cover at non-MPA sites: rescaled to highlight means
Cover during decline (2050)
Minimum cover
Final cover
Caribbean log(scaled cover)
Southwest Pacific log(scaled cover)
Coral Triangle log(scaled cover)
(j)
(k)
(l)
(m)
(n)
(o)
(p)
(q)
(r)
representative
cold
hot
even temperature
pr05
preadapted larvae
eigenvector
evenly spaced
isolation
Strategy

**Appendix S20.** Coral cover at non-MPA sites in the Caribbean (A-C), Southwest Pacific (D-F) and Coral Triangle (G-I) under each strategy scaled by cover when there were no MPAs. Results above the red horizontal line indicate strategies where coral cover was higher than in the simulations with no MPAs while results below the line indicate the opposite. Means are marked as white dots. Panels J-R are rescaled plots of A-I that highlight differences among means.

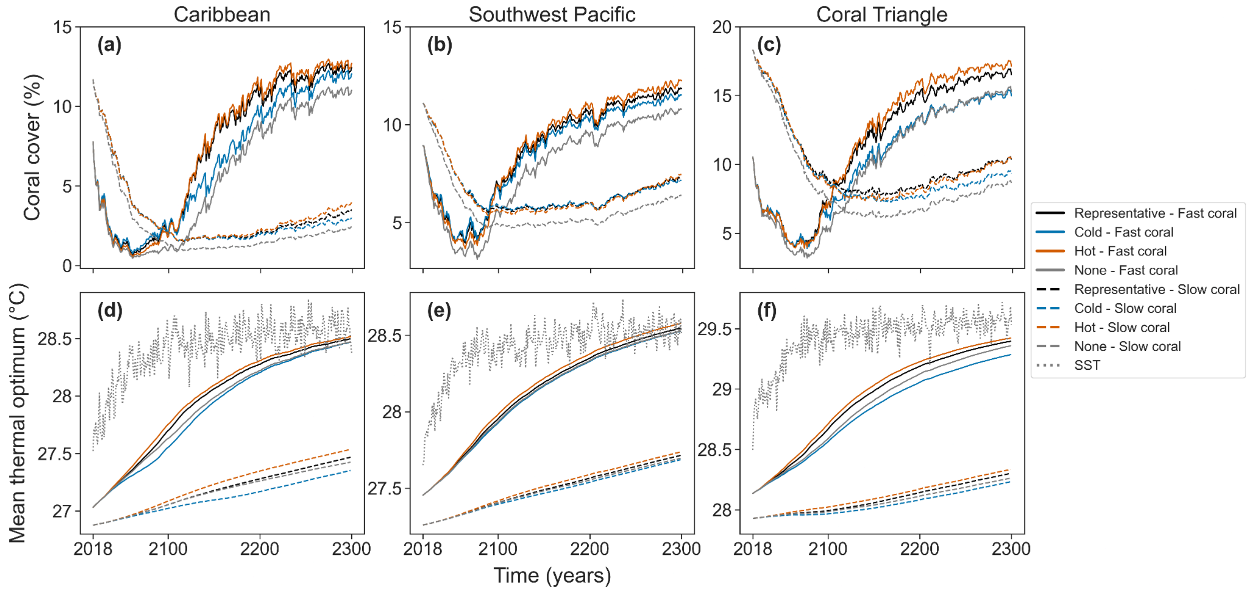


**Appendix S21.** Coral cover (a-c) and mean thermal optimum (d-f) through time for fast (solid lines) and slow (dashed lines). Sea surface temperatures (SST) is shown in dotted gray lines in plots d-f.

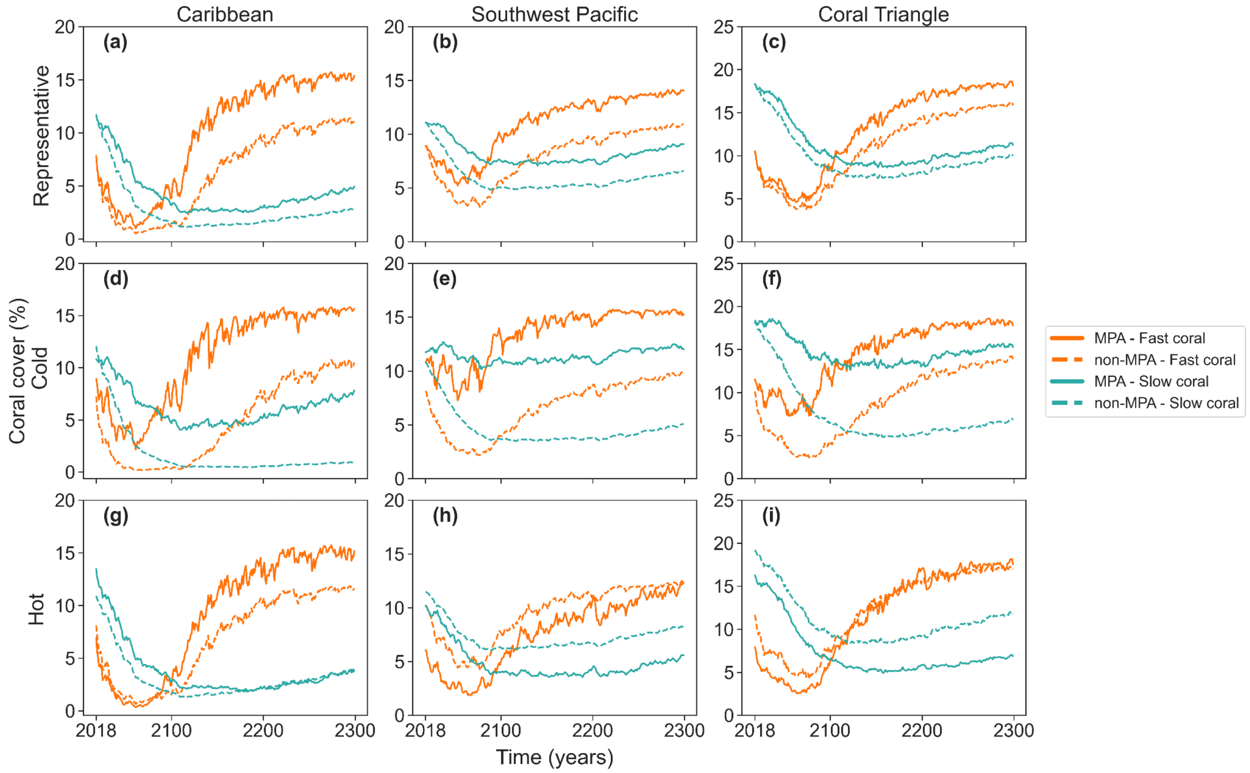


**Appendix S22.** Coral cover through time for the representative (a-c), cold (d-f) and hot (g-i) MPA configurations for reefs inside (solid lines) and outside (dashed lines) MPAs. The fast coral species is shown in orange while the slow coral species is shown in green.

**Appendix S23.** Site characteristics of existing Caribbean MPAs. See Table 1 for definitions.

| **Metric** | **MPA means** | **MPA standard deviation** | **non-MPA mean** | **non-MPA standard deviation** | **Mann-Whitney U statistic** | **Mann-Whitney U p-value** |
|---|---|---|---|---|---|---|
| $T_0$ (°C) | 27.8 | 0.62 | 27.39 | 0.8 | 9625 | 0 |
| *ΔT* (°C) | 1.11 | 0.11 | 1.16 | 0.12 | 10858 | 0.0012 |
| local retention (*L*) | 0.22 | 0.2 | 0.21 | 0.24 | 12074.5 | 0.0361 |
| self-recruitment (*R*) | 0.2 | 0.26 | 0.18 | 0.25 | 11583.5 | 0.0109 |
| destination strength (*S*) | 0.16 | 0.3 | 0.1 | 0.21 | 11206 | 0.0038 |
| pr05 | 0.05 | 0.14 | 0.07 | 0.12 | 10326 | 0.0002 |
| Area ($m^2$) | 6.60E+08 | 6.19E+08 | 6.48E+08 | 7.86E+08 | 12167.5 | 0.0437 |

**Appendix S24.** Site characteristics of existing Southwest Pacific MPAs. See Table 1 for definitions.

| Metric | MPA means | MPA standard deviation | non-MPA mean | non-MPA standard deviation | Mann-Whitney U statistic | Mann-Whitney U p-value |
|---|---|---|---|---|---|---|
| $T_0$ (°C) | 25.9 | 1.24 | 27.95 | 1.42 | 6500.5 | 0 |
| $\Delta T$ (°C) | 0.92 | 0.24 | 0.95 | 0.19 | 20324.5 | 0.0636 |
| local retention (*L)* | 0.38 | 0.39 | 0.31 | 0.36 | 19884 | 0.0342 |
| self-recruitment (*R*) | 0.39 | 0.4 | 0.29 | 0.36 | 18502 | 0.0029 |
| destination strength (*S*) | 0.07 | 0.18 | 0.09 | 0.2 | 20960 | 0.1365 |
| pr05 | 0.36 | 0.37 | 0.13 | 0.27 | 12245 | 0 |
| Area ($m^2$) | 2.16E+02 | 1.05E+03 | 1.76E+02 | 5.01E+02 | 22142 | 0.3824 |

**Appendix S25.** Site characteristics of existing Coral Triangle MPAs. See Table 1 for definitions.

| **Metric** | **MPA means** | **MPA standard deviation** | **non-MPA mean** | **non-MPA standard deviation** | **Mann-Whitney U statistic** | **Mann-Whitney U p-value** |
|---|---|---|---|---|---|---|
| $T_0$ (°C) | 27.35 | 1.31 | 28.75 | 0.61 | 105379 | 0 |
| *ΔT* (°C) | 1 | 0.08 | 1.02 | 0.11 | 205211 | 0 |
| local retention (*L)* | 0.05 | 0.07 | 0.06 | 0.1 | 250444 | 0.14 |
| self-recruitment (*R*) | 0.11 | 0.25 | 0.08 | 0.17 | 254420.5 | 0.253 |
| destination strength (*S*) | 0.32 | 0.3 | 0.28 | 0.23 | 258362 | 0.402 |
| pr05 | 0.16 | 0.23 | 0.08 | 0.17 | 185071 | 0 |
| Area ($m^2$) | 3.42E+08 | 3.84E+08 | 1.31E+08 | 1.69E+08 | 153323 | 0 |

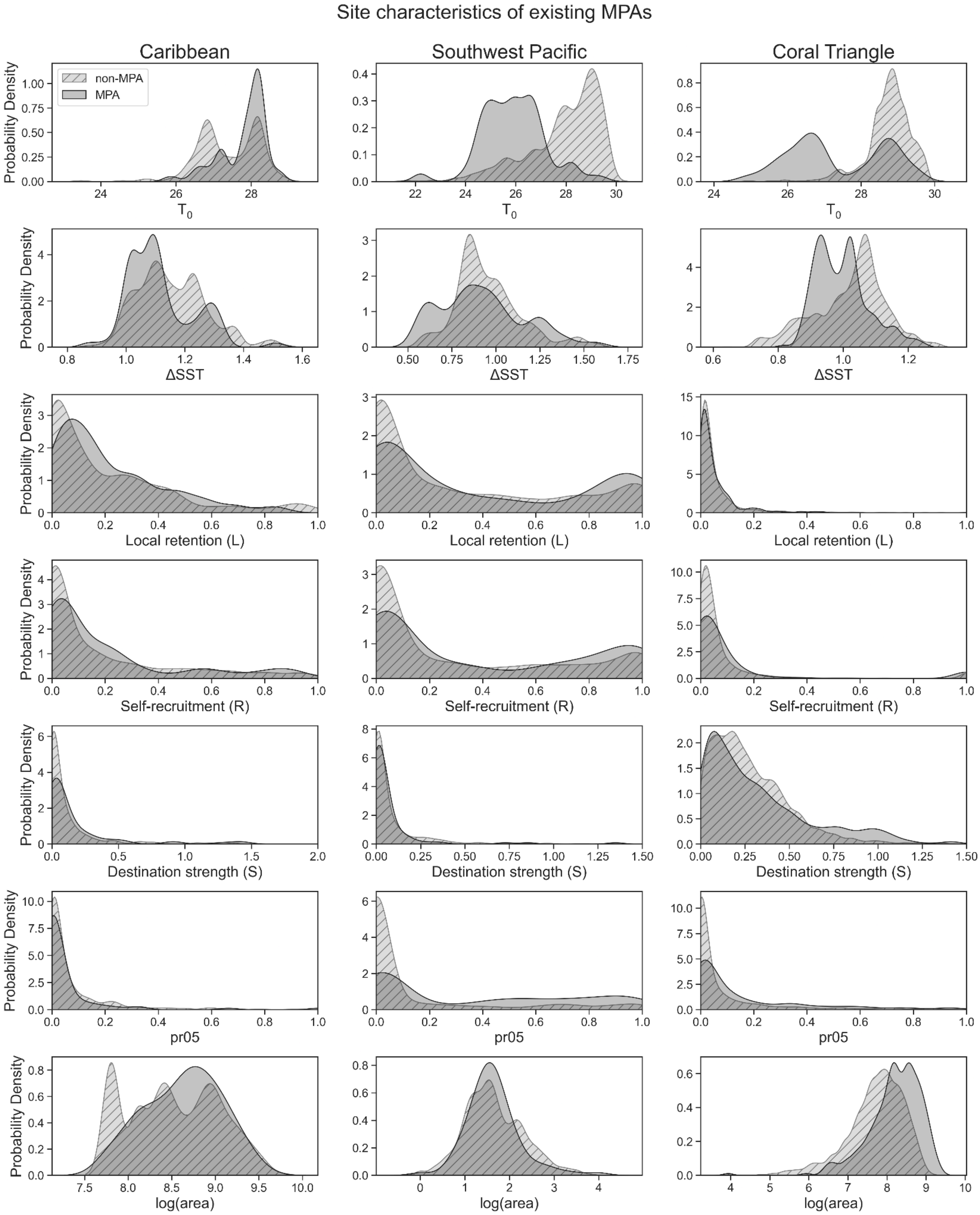
Site characteristics of existing MPAs
Caribbean
Southwest Pacific
Coral Triangle
non-MPA
MPA
Probability Density
$T_0$
ΔSST
Local retention (L)
Self-recruitment (R)
Destination strength (S)
pr05
log(area)

**Appendix S26.** Kernel density functions for MPA and non-MPA sites across all regions (columns) of the following site characteristics (rows, top to bottom): initial sea surface temperature ($T_0$), change in sea surface temperature ($\Delta T$), local retention ($L$), self-recruitment ($R$), destination strength ($S$), pr05 and log(area).

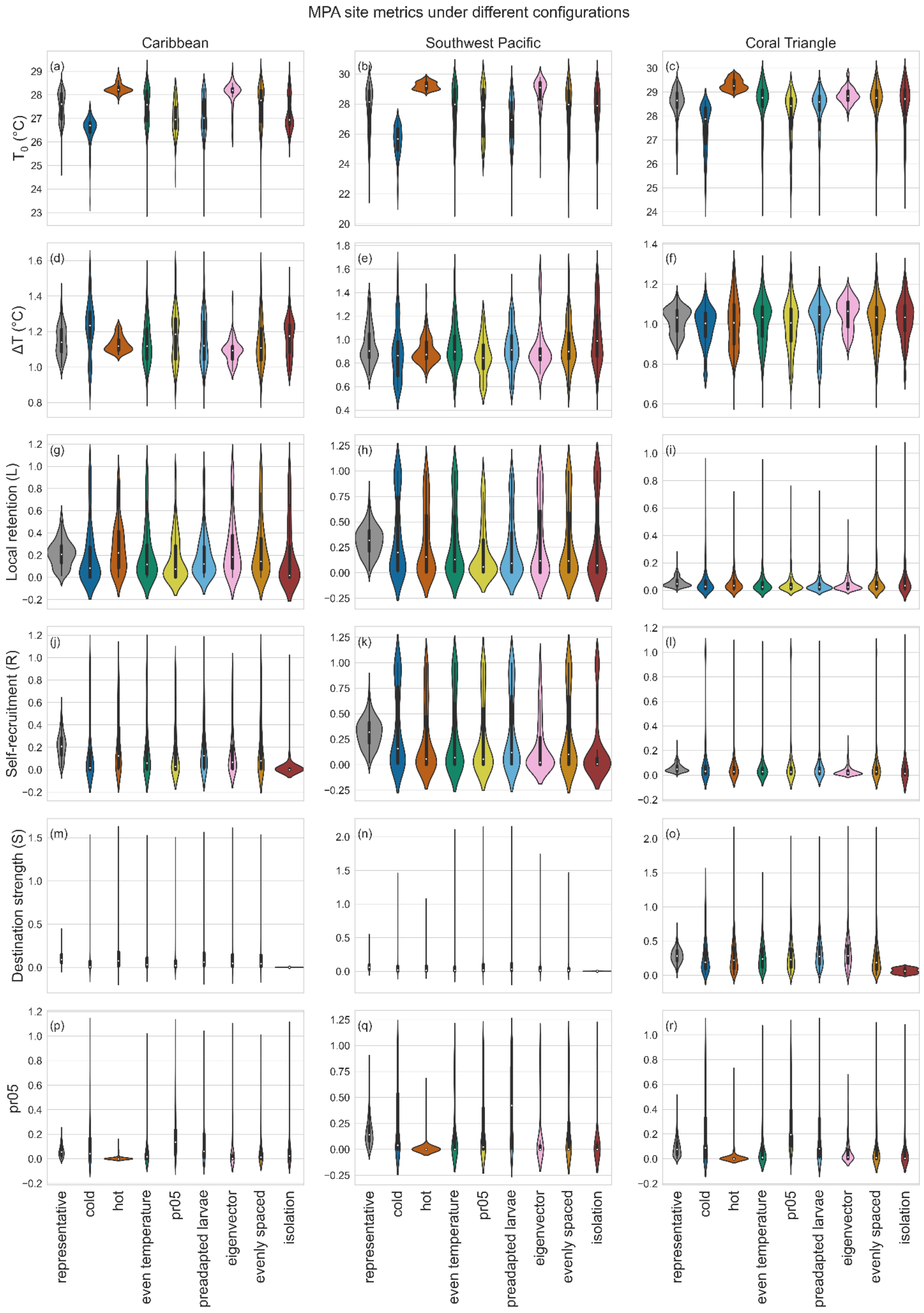
MPA site metrics under different configurations
Caribbean
Southwest Pacific
Coral Triangle
$T_0$ (°C)
ΔT (°C)
Local retention (L)
Self-recruitment (R)
Destination strength (S)
pr05
(a)
(b)
(c)
(d)
(e)
(f)
(g)
(h)
(i)
(j)
(k)
(l)
(m)
(n)
(o)
(p)
(q)
(r)
representative
cold
hot
even temperature
pr05
preadapted larvae
eigenvector
evenly spaced
isolation

**Appendix S27.** Violin plots comparing initial sea surface temperature $T_0$ (A-C), change in sea surface temperature *ΔT* (D-F), local retention *L* (G-I), self-recruitment *R* (J-L), destination strength *S* (M-O) and pr05 (G-I) of sites selected for MPAs across the three regions.
